# Ecological scaffolding and the evolution of individuality: the transition from cells to multicellular life

**DOI:** 10.1101/656660

**Authors:** Andrew J Black, Pierrick Bourrat, Paul B Rainey

## Abstract

Evolutionary transitions in individuality are central to the emergence of biological complexity. Recent experiments provide glimpses of processes underpinning the transition from single cells to multicellular life and draw attention to the critical role of ecology. Here we emphasise this ecological dimension and argue that its current absence from theoretical frameworks hampers development of general explanatory solutions. Using mechanistic mathematical models, we show how a minimal ecological structure comprised of patchily distributed resources and between patch dispersal can scaffold Darwinian-like properties on collectives of cells. This scaffolding causes cells to participate directly in the process of evolution by natural selection as if they were members of multicellular collectives, with collectives participating in a death-birth process arising from the interplay between the timing of dispersal events and the rate of resource utilisation by cells. When this timescale is sufficiently long and new collectives are founded by single cells, collectives experience conditions that favour evolution of a reproductive division of labour. Together our simple model makes explicit key events in the major evolutionary transition to multicellularity. It also makes predictions concerning the life history of certain pathogens and serves as an ecological recipe for experimental realisation of evolutionary transitions.

## INTRODUCTION

Evolutionary transitions in individuality (ETIs) are central to the emergence of biological complexity^1–3^. Each ETI involved the formation of collective-level entities from the interaction of particles^4,5^. For example, chromosomes evolved from the joining of once independently replicating genes. Multicellular life evolved from independently replicating cells. In certain insects, eusociality evolved from independently replicating multicellular types.

Central to each of these transitions was the emergence of properties at the newly formed level that allowed individuals — at the newly formed level — to participate directly in the process of evolution by natural selection^5–9^. This required newly formed collectives to be discrete and vary one to another, to reproduce and to leave offspring that resemble parental types^10^. These essential and intertwined Darwinian properties of variation, differential reproduction and heredity are such fundamental features of living systems that it is easy to overlook the fact that individuality is a derived state and in need of evolutionary explanation^3,7–9,11–13^.

With focus on multicellular life, it is evident that reproduction, in even simple multicellular forms, is a complex process^9,11,12,14^. It is therefore tempting to invoke selection as its cause. But this is problematic because the earliest collectives lacked capacity for collective-level reproduction and thus to invoke selection at the collective level as the cause of collective-level reproduction is to invoke the trait requiring explanation as the cause of its own evolution. Clearly such an explanation is unsatisfactory.

One way to avoid this dilemma is to recognise opportunities for co-option of pre-existing cellular traits. For example, in the colonial volvocine algae, group formation evolved by co-option and expansion of cell cycle regulation evident in unicellular *Chlamydomonas*^15^. In experimentally evolved snowflake yeast, collective-level reproduction emerged via co-option of apoptotic capacity already apparent in single cell precursors^16^.

We do not wish to downplay the importance of co-option, but there is conceivable value in asking whether Darwinian properties at the collective level might emerge in the absence of co-option. Such a take-nothing-for-granted line of inquiry presents a challenge as it requires conceiving possibilities for the emergence of properties essential for collectives to participate in the process of evolution by natural selection from starting states that lack any manifestation of collective-level Darwinian properties. In essence it begs explanations for how Darwinian properties might emerge from non-Darwinian entities and therefore by non-Darwinian means. Solutions stand to inform not only how multicellular states arise from single cells, but how Darwinian properties might emerge during each of the major evolutionary transitions, including that from non-living matter.

A solution that we advance draws heavily on ecology, the significance of which we suggest has been overlooked — even though the importance of population structure has been emphasised by literature on the levels of selection^17^. It recognises that Darwinian properties can be “scaffolded” by the environment — that these properties can be exogenously imposed in such a way as to cause lower level entities (e.g., cells) to become unwitting participants in a selective process that occurs over a longer timescale than the timescale over which cell-level selection occurs, and as part of a larger entity (a collective). In time, such exogenously imposed — Darwinian-like — properties stand to become endogenous features of evolving systems (for development of general views on scaffolding processes see^18^).

Ecological scaffolding underpinned a recent (and on-going) experimental exploration of the evolution of multicellularity^19,20^. Discrete lineages established from the bacterium *Pseudomonas fluorescens* were propagated under conditions that required, for long-term persistence, repeated completion a two-phase life cycle involving soma and germ-like states. In the experiment, variation was discretised using glass microcosms, but the design is loosely analogous with an environment such as a pond in which reeds extend from the water^19,21^. Each reed allows establishment of a single microbial mat (the soma-like phase), with the spacing of reeds ensuring variation at the level of mats. Mats that collapse, for example, through physical disturbance, are extinguished, allowing the possibility that an extant mat might, via production of a dispersing (germ-like) phase, increase its representation among the population of mats. Thus, the possibility of a selective process unfolds at the level of mats. After ten lifecycle generations, the fitness of derived mats significantly improved, with the most successful lineage having even evolved a simple genetic switch that ensured reliable developmental change between soma and germ-line phases^19^. Not only does this study demonstrate that scaffolding works, but it also shows that externally imposed Darwinian properties can begin the shift toward endogenisation exemplifying Van Valen’s view that “*evolution is the control of development by ecology*”^22^.

Our goal here is to show how a minimal set of ecological conditions (and ensuing evolutionary responses) can effect evolutionary transitions in individuality. We take inspiration from the the experimental *Pseudomonas* studies, but simplify the ecological context in order to produce a minimal mechanistic model. Although our focus is the transition from single cells to multicellular life, we argue that the concept of ecological scaffolding is relevant to other transitions. It also makes predictions concerning the life history of certain pathogens and serves as an ecological recipe for top-down engineering of populations and communities.

## RESULTS

### Scaffolding Darwinian properties

Before moving to mathematical models, we describe the simplest conceivable example of a population structure that confers Darwinian-like properties on collectives of particles (cells). Consider an environment in which resources are distributed across patches. A single cell founds a patch. Available resources allow exponential growth of the founding type, however, because resources within patches are finite, they are rapidly depleted causing the population to decline (equations describing the birth / death process and relationship with resources are described in the Methods section). Long-term persistence of cells requires dispersal to a new patch. Dispersal occurs at a fixed regular time interval via, for example, some external factor such as wind, water splash or tidal flow.

Cell fate within the environment of patches depends on performance over two timescales. The first timescale is defined by the doubling time of cells. The second is defined by the timing of dispersal events. To make apparent impact of the second timescale on ensuing evolutionary dynamics, consider a second variant cell. This type grows faster than the former (cells consume resources more rapidly), which means that in a patch founded by both types, faster growing cells rapidly exclude slower growing cells. In the following we therefore limit the number of colonisers to a single founding cell type, thus limiting within patch competition.

Consider a single slow (depicted in Figure 1 as green) and fast (blue) cell that colonise separate patches (Figure 1). Cells of both types grow and divide, but different growth rates mean that blue cells deplete resources more rapidly than green cells. If dispersal occurs early (Figure 1A) when cells are in exponential growth, then the number of extant blue cells exceeds the number of green cells and thus future recursions of patches are dominated by blue cells (Figure 1B). If, however, dispersal occurs at a later time point, for example, once resources are depleted and population size is in decline, as in Figure 1C, then future patch recursions are dominated by green types despite the fact that within a patch, green types lose in competition with blue types (Figure 1D).

**Figure 1.**
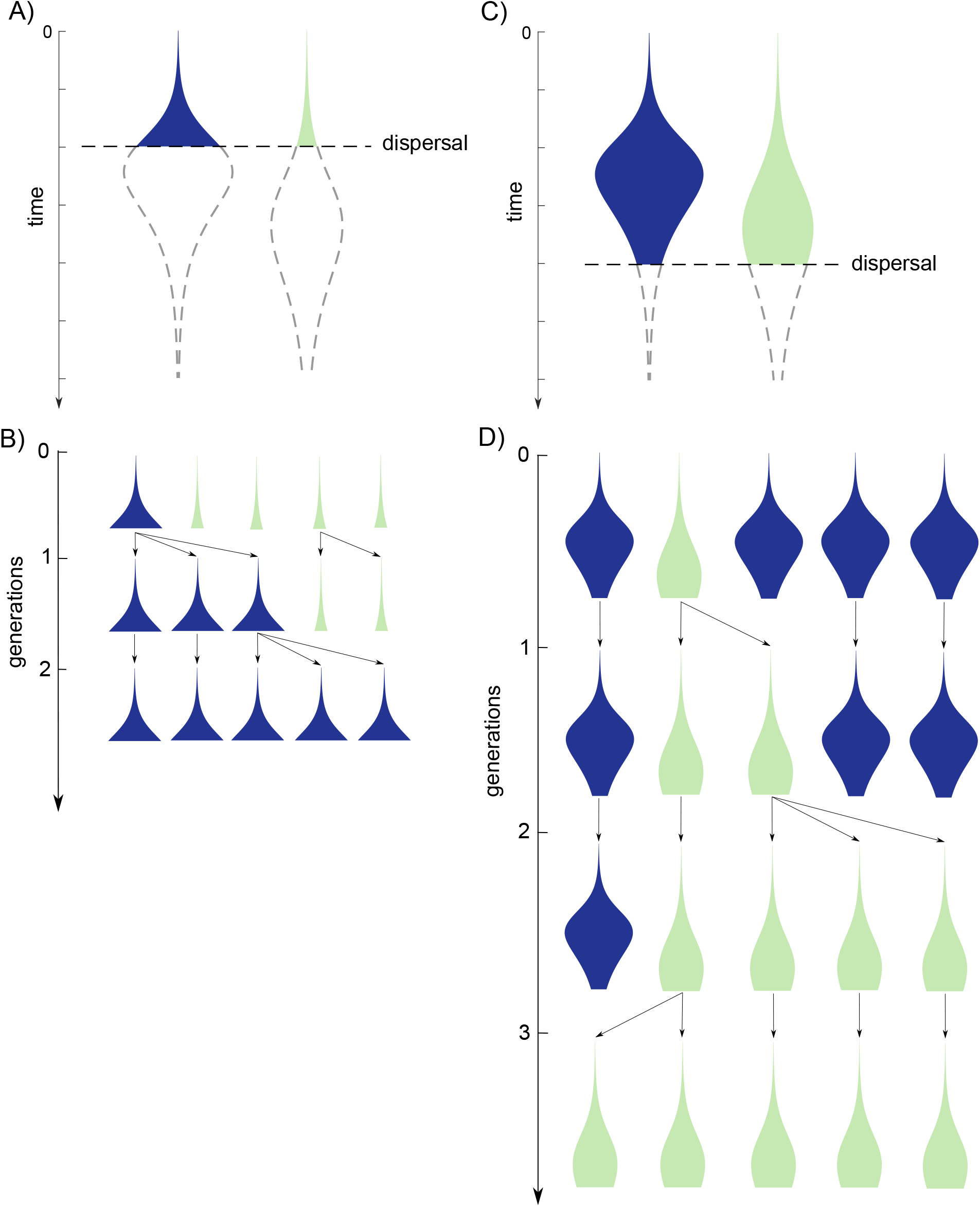
Scaffolding Darwinian properties. Patchily distributed resources provide opportunity for two cell types (blue and green) to replicate (blue cells grow faster than green according to equation 1 (Box). Single cells of each type colonise discrete patches at time t = 0 and consume resources. Difference in growth rate means the relationship between cell density and time differs for blue versus green populations. Should a dispersal event occur during exponential growth (dashed line) then more blue cells will be dispersed relative to green (A) and thus the blue population will be more successful over the long term (B). Conversely, should dispersal occur at a later stage and after resources are depleted (C), then the population of green cells will out compete green over the long term (D).

Thus far, our focus has been the consequences of this population structure on the long-term fate of cells with different growth rates, but it is possible to switch perspective: there is a coupled evolutionary dynamic occurring at the level of patches (Figures 1C and 1D). Patches manifest Darwinian-like properties of variation (spatially distributed resources ensure that variation is discretised and that patches vary one to another), differential reproduction (successful patches give rise to patches via dispersal) and heredity (offspring patches are likely to resemble parental patches because new patches are founded by single cells) that are also features of the founding cells. These properties are externally imposed (scaffolded) on patches by virtue of the structure of the environment.

Note that we refer to the properties of patches as “Darwinian-*like*”. Indeed, it makes no sense to think of patches as multicellular organisms (they are not) — if the ecological scaffold was to be removed (patchily distributed resources and a means of dispersal) — the Darwinian-like properties of the patches would instantly disappear. Yet, under the scenario outlined, cell fate is determined by selective conditions operating over the second (longer) timescale, just as if the cells themselves were members of multicellular collectives. Such a scaffolded framework of patch-level selection, based on nothing other than patchily distributed resources and a means of dispersal between patches, establishes conditions sufficient for the evolution of traits that are adaptive at the level of patches. We elaborate the mechanistic bases using models developed in the following section, but firstly comment on the connection between the heuristic model outlined above and previous models.

The basic structure of our model, with patchy environments and a dispersal process, bears some similarity to various models of group selection, including models of Wright’s shifting balance theory^23–25^, Maynard Smith’s haystack model^26^ and others^27,28^. Our model differs from these in both its emphasis on mechanism and what it attempts to explain. Earlier models, with some exceptions^29,30^, are phenomenological, that is, they are constructed in a way that captures the relationship between variables and the problem under consideration^31^. Hence, while they capture the action of various forms of group selection, they are compatible with a range of causal structures. In contrast, our model is mechanistic and so evolutionary change can be understood in terms of the underlying non-linear dynamics of particles and the feedback that arises from interaction with the timing of dispersal. This allows transparent elaboration of the model, including the possibility of future systematic explorations of the robustness of ecological scaffolding. Another key difference to earlier standard group selection models is that our focus is the evolution of groups as units of selection in their own right and not the effect of group structure on the evolution of behaviours that are costly to individual cells, such as altruism^27,32^.

Further connections though are possible and we draw particular attention to connections with the levels of selection literature^3,33^. Early stages of evolutionary transitions are often seen as being encapsulated by the multilevel selection 1 (MLS1) framework, as defined by, for example, trait group models^27^. In these models group fitness is the average (or sum) of the fitness of the cells that comprise collectives. The transition completes once collectives become individuals and units of selection in their own right. At this point fitness of collectives is defined independently of cell fitness and relevant models fall within the multilevel selection 2 (MLS2) framework^4^. The shift between levels — involving transference of fitness from cells to collectives — has been difficult to capture^3^. An important insight came from Michod and colleagues^4,34,35^ who articulated and modelled the concept of fitness decoupling: the notion that the shift between MLS1 and MLS2 involves collective level fitness “decoupling” from lower level (cell) fitness. The heuristic model depicted in Figure 1 and elaborated below captures both MLS1 and MLS2 dynamics in a single model with the transition between the two levels arising as an emergent feature.

### Evolution in nested Darwinian populations

To explore the evolutionary dynamics of the above ecological model we allow mutation to affect the growth rate of individual cells. With such a model it becomes possible to determine the effect of the timing of dispersal — the second timescale — on the dynamics of within- and between-patch competition. Mathematical details are provided in the Methods and Supplementary Information file.

The full evolutionary model consists of *M* patches that are each founded by a single cell of a single phenotype (growth rate *β*). Cells within patches replicate and consume resources with mutation giving rise to types that vary in growth rate. Once resources are depleted the population size within patches declines. After a fixed time interval, *T*, which defines the second timescale, dispersal takes place. Dispersal is effected by randomly selecting *M* patches (with replacement) in proportion to the number of cells within each patch, and then randomly selecting a cell, within chosen patches, in proportion to numbers within the patch. In effect, the procedure is equivalent to pooling all viable cells from all patches at the time of dispersal and picking *M* cells at random in a manner akin to an extreme form of a trait group model^27^. The dispersal regime thus rewards patches containing the greatest number of cells.

The bottleneck wrought at the moment of dispersal means that types founding new patches are freed from competition with faster growing types. Figure 2 shows the number of cells within a patch for a single realisation with initial (arbitrarily chosen) growth rate *β* = 1.8. The bottleneck imposes a strong homogeneity on the composition of the patch as the original population has to grow significantly before mutants start to arise. The peak (maximum) number of cells within the patch is reached at time *T* = 16.5, thus for cells with this initial growth rate, setting a dispersal time of *T* = 10 is fast (i.e., still within the exponentially growing phase), and *T* = 30 is slow (cells have significantly declined since their peak numbers).

**Figure 2.**
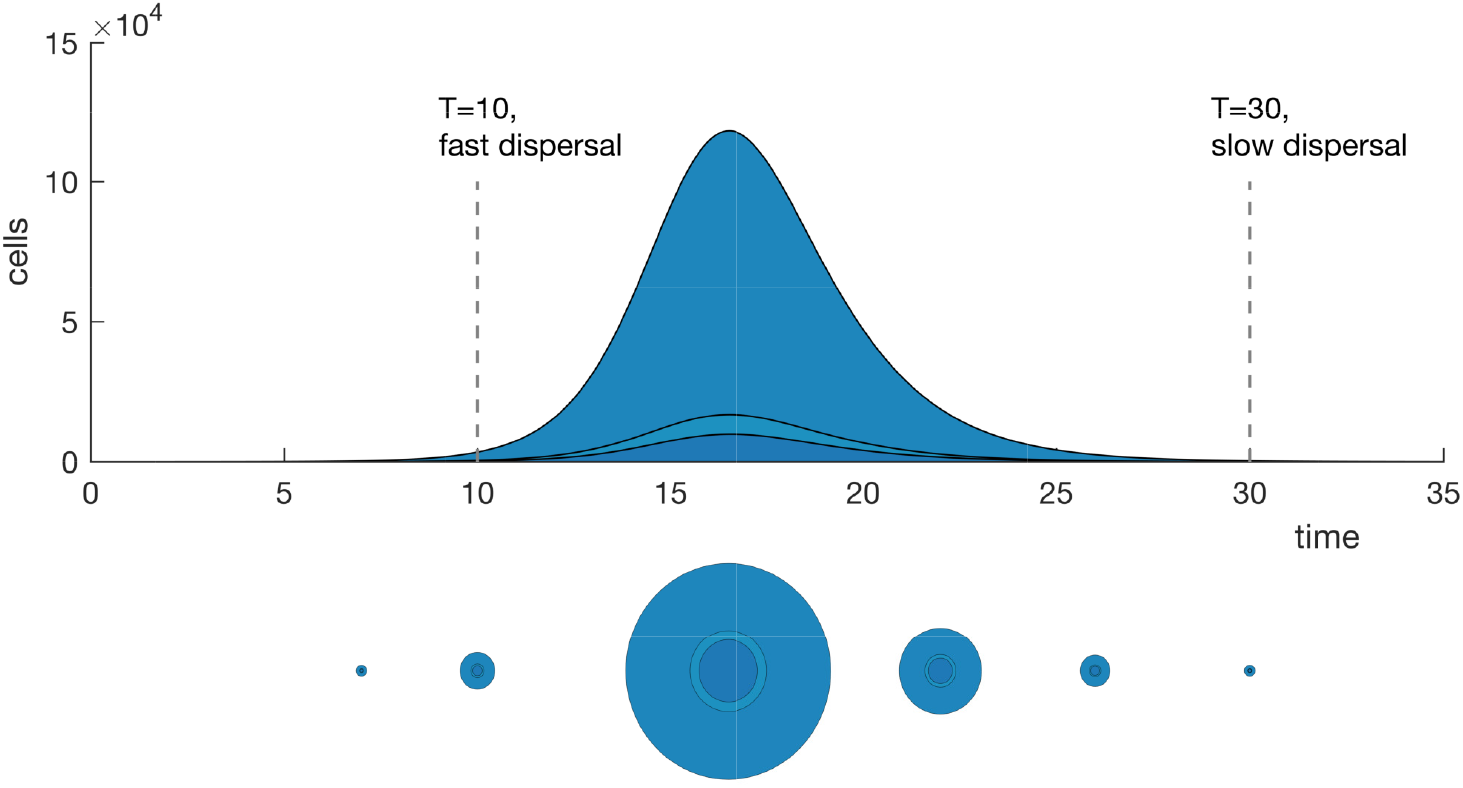
Single realisation of the within patch model with mutation. The number of cells is plotted as a function of time starting from a single cell with growth rate *β* = 1.8 and the amount of resource *N*=10^6^. The darker shaded regions show the numbers of mutant cells. Darker and lighter colours correspond to faster and slower growing cells, respectively. The circles below the main plot are representations of cell numbers (used in the video and Figures 3 and 4) at times 7, 10, 16.5, 22, 26 and 30. In this representation, the *area* of each region is proportional to the number of cells of each type within the patch (with the same colour scheme). The peak number of cells within the patch is reached at time ~16.5, thus for the initial growth rate of *β* = 1.8, setting a dispersal time of T=10 is considered fast, and T=30 slow (shown by the dashed lines).

**Figure 3.**
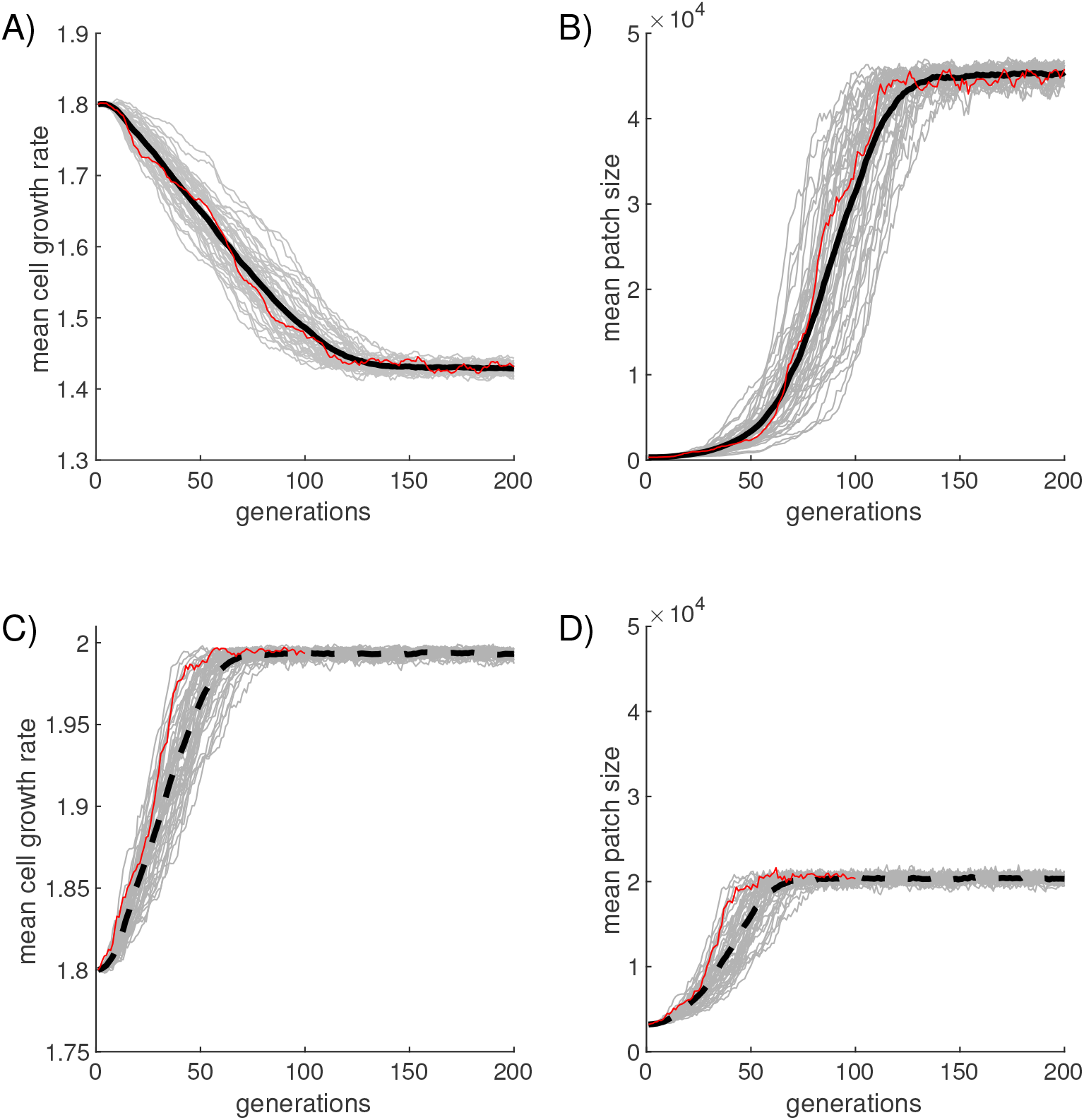
Effect of dispersal timescale on properties of cells and patches. Each grey line is from an independent realisation of the stochastic model. The solid black lines are averages over 50 realisations. A and B show the evolution of the average growth rate over all patches in the generation with slow dispersal (*T*=30). C and D are the same but with fast dispersal (*T*=10). Both regimes start with a homogenous population of cells with *β* = 1.8 and in both cases the average patch size increases, but for slow dispersal this is achieved by cells decreasing their average growth rate. Figure 4 shows the 64 patches at the moment of dispersal for two single realisations (shown in red above) after a number of generations.

**Figure 4.**
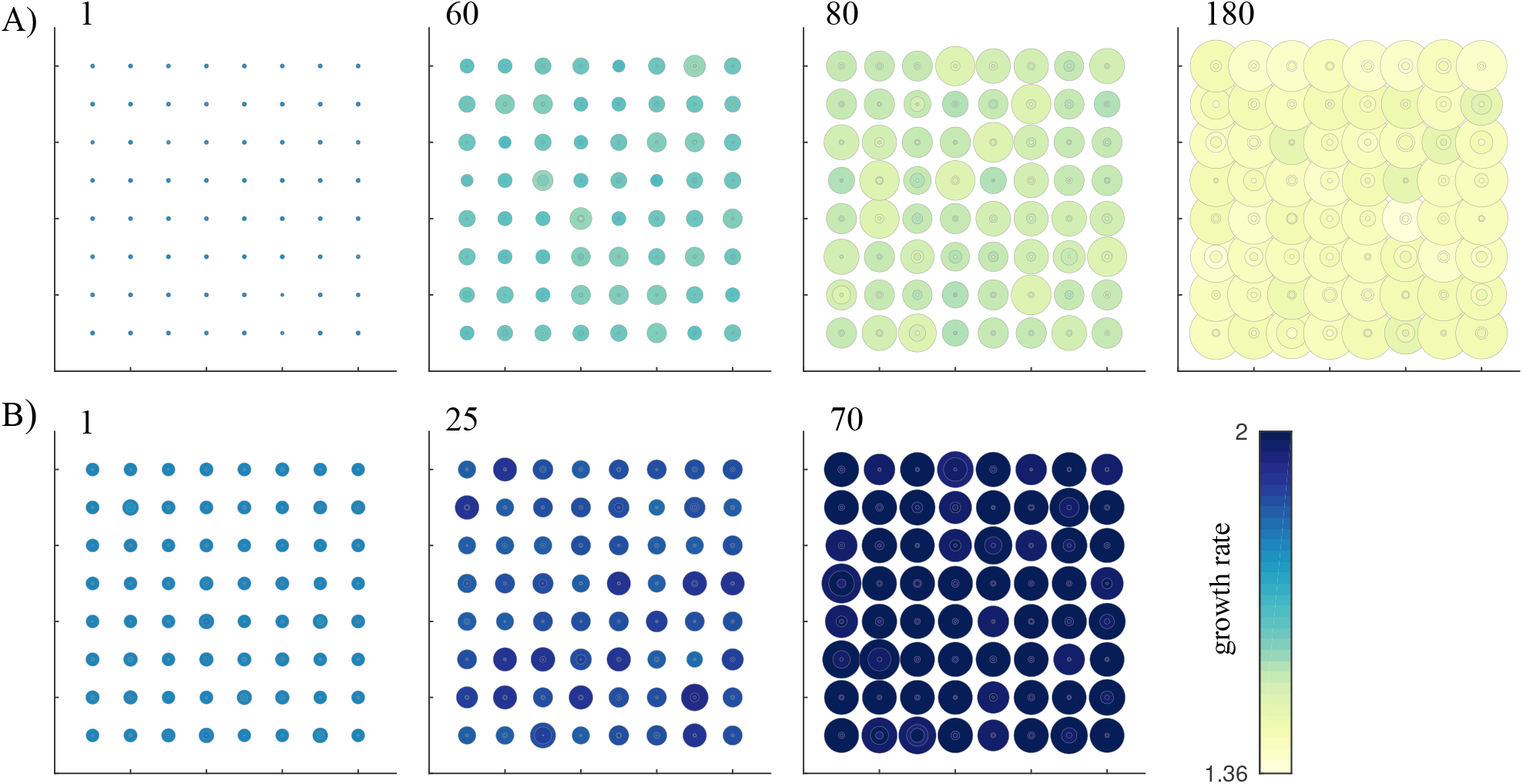
Evolution of patch size under slow and fast regimes. The dynamic of patch-size evolution and corresponding effects on cell growth rate under slow (A) and fast (B) dispersal regimes. Movies of simulations are shown in Supplementary Movie Files 1 and 2. Colour corresponds to cell growth rate and patch size is proportional to the number of cells within patches at the moment immediately prior to dispersal. Generation number is indicated above each panel and corresponds to the realisations highlighted in red in Figure 3.

From the patch perspective, the bottleneck reduces within patch variation and ensures high fidelity of transmission of patch phenotype (the size of the patch at the time of dispersal). Cells chosen for dispersal are individually transferred to new patches thus marking the founding of a new generation of patches. The mechanistic nature of the model allows the average growth rate of cells within a generation, number of cells in patches at the time of dispersal, and genealogy to be determined.

Figure 3 shows the time resolved dynamics of 50 independent realisations of the full evolutionary model, where patches experience 200 recursions, under slow (*T* = 30) and fast (*T* = 10) dispersal regimes. In these simulations the maximum cell growth is set to rate of *β* = 2, which in a real system would arise from chemical and physical constraints to the rate of cell replication. The state of populations at the time of dispersal are shown in Figures 4A and 4B. Single realisations of the model under slow and fast dispersal regimes are also shown in Supplementary Movie Files 1 and 2.

Under both fast and slow dispersal regimes patch fitness (the number of cells within patches at time of dispersal) increases rapidly before reaching a plateau (Figure 3B and 3D). This is consistent with predictions arising from the logic of Darwinism: imposition (by ecological scaffolding) of Darwinian-like properties on patches ensures patches participate in a selection process akin to evolution by natural selection, one that could be the starting point for patches to be units in their own right, provided they eventually acquire features classically associated with evolutionary individuals^5,36^. The plateau arises because under the slow dispersal regime growth rate evolves to maximise the number of particles available at the time of dispersal. Under the fast dispersal regime, the plateau is a consequence of reaching the maximum limit imposed by the growth rate. As this maximum rate is arbitrarily set, allowing it to increase would result in the evolution of patches of larger final size.

The cause of enhanced evolutionary success of patches resides in properties of individual cells. Under both fast and slow dispersal regimes, selection favours patches that harbour the greatest number of cells at the time of dispersal. Under both regimes fast growing cells outcompete slow growing types within patches, however, under the slow dispersal regime, selection rewards patches containing slow growing mutants and selects against patches dominated by fast growing cells. The opposite is true of patches evolving under the fast dispersal regime.

Under the slow dispersal regime this results in the seemingly counter intuitive finding that patch fitness increases at the expense of cell fitness (Figures 3A and 3B). Yet within our model, this is readily explained: fitness of a cell is measured over the short timescale while patch fitness is measured over the long timescale^37,38^. This captures precisely — and explains mechanistically — the notion of “fitness decoupling” thought to occur during the earliest stages of the evolution of multicellular life, but which has often been difficult to intuit^3,39^.

Under the fast dispersal regime, fast growing cells are favoured both within patches and over the second timescale. From the perspective of the evolution of multicellular life, the selection regime imposes the same directionality at both timescales leading to the view that fitness at both timescales (levels) are “coupled”.

It is interesting to note the difference in speed of the selective response under the two dispersal regimes and also the magnitude of difference in patch population size at equilibrium. The slower response under the slow dispersal regime is a consequence of the time taken for slow growing mutants to invade from rare in the face of within-patch competition for fast growth. The maximum population size under the fast dispersal regime (for these parameters) is a consequence of the imposed maximum limit on the growth rate. If faster rates were allowed, then larger population sizes would evolve (up to a limit where rate maximises patch size at the dispersal time).

Supplementary Figures S3A and S3B show the evolutionary fate (genealogy) of 10 independent lineages under the slow and fast dispersal regimes, respectively. Mapped on the phylogenies are changes in cell growth rate and patch size at time of dispersal. That a genealogical representation is possible derives from both the mechanistic nature of the model and the fact that patches are founded from single cell types. Movie versions of the Figures S3A and S3B are shown in Supplementary Movie Files 3 and 4.

It is important to emphasise that the parameters and timescales chosen in the above simulations are arbitrary. For any initial growth rate > 1, it is always possible to choose fast and slow dispersal times relative to the time of the peak patch population that will result in selection over the timescale of dispersal feeding back to affect the growth rate of cells. In contrast, if the dispersal time is equal to the initial peak time, then no evolutionary change in cell growth rate will be observed.

### Synchronicity of dispersal

The analysis above assumes a predictable environment: resources within patches are identical for all patches, the time at which patches are seeded by new colonists is fixed, and so is the time interval until dispersal. In this section we explore relaxation of this strict ecology and do so by two adjustments to the model. In the first, variance in the time at which patches are seeded with new propagules is introduced. In the second, the level of resource available in each patch is varied.

Variance in the growth time until dispersal is introduced by allowing each patch to grow for a period of time *τ* ∼ *norm*(*T*, *σ*_*T*_). This leads to variance in the size of the population of cells within patches at the time of dispersal. We consider only mean dispersal times that result in tension between the short and long-term interest of cells and patches.

Figure 5 shows evolution of the average growth rate of cells and patch size for a fixed mean (*T* = 30), but with increasing variance in the growth time before dispersal. For small values of *σ*_*T*_ the dynamics show little change, but the effect is pronounced for larger values of *σ*_*T*_. The average growth rate at equilibrium increases slightly, but the main effect is reduction in the average time for the equilibrium state to be realised. The results are readily explained: increasing levels of variance reduces correlation between growth rate of the initial founding cell and the phenotype of the patch (its size at dispersal). Hence the correlation between the size of a patch at dispersal and the size of its parent is reduced, thus reducing the relationship between parent and offspring patches.

**Figure 5.**
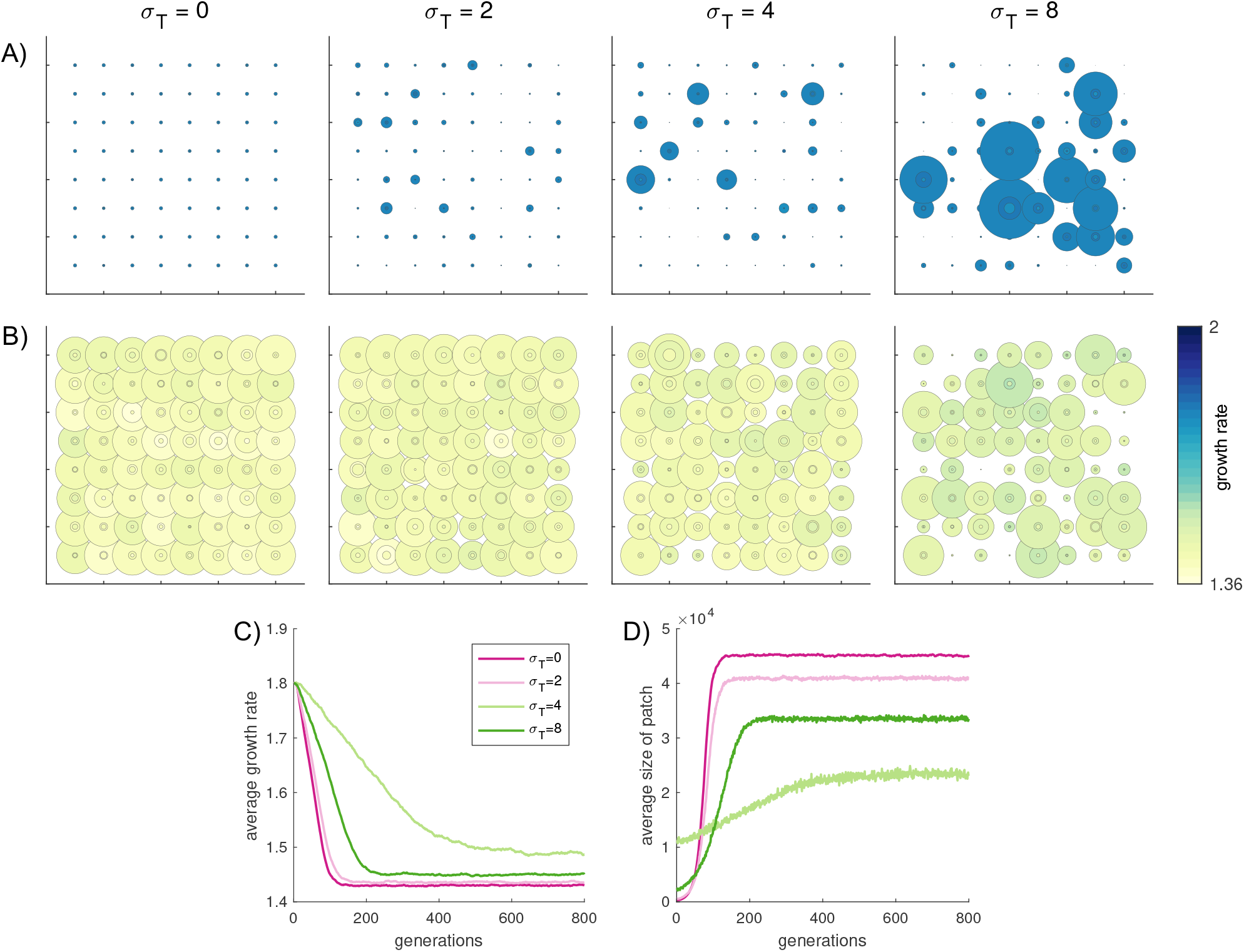
The effect of variance in the growth times within a generation on the evolutionary dynamics. The top panels show the size and composition of patches for single realisations of the first generation (A) and a generation once equilibrium has been reached (B) respectively for increasing *σ*_*T*_. Panels (C) and (D) show the evolution in the average growth rate and patch size over 800 generations respectively from 50 realisations of the model. All simulations start from ell with a homogeneous growth rate of 1.8.

Differences in average patch size, both at the beginning and end of the process are explained by distributions in patch size that are induced by the variance in growth period across the patches in a single generation. This is visualised in Figure 5A and B. With no variance (*σ*_*T*_ = 0), the size of patches is homogeneous, with small size differences caused only by mutations within the patch. Variability in growth time leads directly to variability in patch size at dispersal time. As variance in *τ* increases so does variance in patch size.

The system begins out of equilibrium, that is, with dispersal time being much longer than the average time for populations within patches to peak. As variance in growth time increases, the likelihood that patches with large numbers of individuals at the time of dispersal increases, thus skewing the size distribution of patches in the next generation of patches (Figure 5A). Once at equilibrium (Figure 5B), the average peak population time within a patch coincides approximately with the mean growth time before dispersal. This means that patches in which cells grow for shorter or longer times harbour populations with reduced population sizes compared to the average, hence mean patch population size at equilibrium is reduced when compared with the treatment in which populations are seeded at the same time. Also affected by increasing variance is average cell growth rate. This decreases even when the time of patch seeding shows maximal variance, but both the rate of reduction is lowered and the equilibrium growth rate is higher. Thus as long as average patch size correlates with the growth rate of the initial colonist cell, evolution in growth rate is observed.

The second route to reduce environmental predictability is via adjustments in the initial amount of resource in each patch. The simplest way is to introduce variance in *N*, the initial resource in a patch. Interestingly, this has minimal effect on the evolutionary dynamics (see Figure S4): individual realisations become noisier, but the average rate of change in the growth rate and the equilibrium remain unchanged. This is because variability in *N* introduces variability in the number of cells within each patch (patch size) at the time of dispersal (more resource allows more growth), but the time for the population to peak is independent of *N* due to scaling of the growth rate assumed in the model (more resource implies a larger patch volume so the concentration per unit volume remains the same). This type of variability therefore creates variation in patch population size that is symmetric about the average, hence mean patch size and the mean dynamics do not change. This is in contrast to the situation described above where variability is introduced into the seeding time, which leads to non-symmetric patch size distributions.

An alternate manipulation is to vary the concentration of resource in each patch, while keeping the volume fixed (see Methods). This introduces differences in the time for each patch population to reach its peak, leading to effects similar to those observed for changes in the timing of patch seeding (Figure S5). For this case there is a limit to the degree of variance that can be added because if the concentration of resource is too small, the population fails to grow.

Overall these results show that scaffolding of Darwinian properties on collectives is robust to variance in the timing of dispersal: despite introduction of significant variation in the time of seeding patches and concentration of resources within patches, cells within patches continue to evolve as if members of multicellular collectives, with the fitness of cells decoupling from the fitness of patches. Evident in the analysis is the importance of the interaction between timescales. The time at which the population peaks and its correlation with average patch size as determined by the initial growth rate and resource, determines the speed of the evolutionary dynamics and its equilibrium state. Analysis using a Price equation-based approach^40^ would likely lead to similar conclusions, but this would be difficult to derive and calculate due to the non-linear nature of our model. Additionally, without an underlying mechanistic model, the Price approach would not provide an explanation for the observed dynamics or the variability in individual trajectories.

### Evolution of patch traits

The above model shows how a second timescale, defined by dispersal events necessary for establishment of new patches, affects the evolution of cell growth rate, and how changes in cell growth rate affect the evolutionary dynamics of patches. From the patch perspective, derived patches are more fit than ancestral patches, but this is not a consequence of traits adaptive at the patch level. Under both slow and fast dispersal regimes, selection favours cells whose growth rate maximises the number of cells available for dispersal. Changes in cell growth rate thus fully explain the evolutionary dynamics of patches. This cell-level perspective further emphasises the previous comment that patches are not to be confused with even the most basic manifestations of multicellular life forms.

Nonetheless, an ecological scaffold that couples short and long-term timescale dynamics establishes kin groups^41^, conducive to the evolution of traits adaptive at the level of patches. By this we mean traits that would be difficult to explain from the view point of cells. This prediction becomes intuitive upon switching perspectives, from cell-level to patch-level. Although patches are endowed with Darwinian-like properties, there is scope for patches to evolve genuine Darwinian properties — in a ratchet-like manner^42^ — so that patches participate in the process of evolution by natural selection and thus become bearers of adaptations at the patch level.

What might such patch-level adaptations entail and what might constitute their mechanistic (cell-level) basis? A fundamental requirement given the need for patches to pass through single bottlenecks at each recursion is evolution of a stochastic epigenetic switch (a simple developmental programme), such as observed previously in numerous experiments^43,44^ including those arising from experimental explorations of the evolution of multicellular life^19^.

To investigate this possibility the basic model is extended to include two types of cell. The first type, which we denote *G*, is essentially the same as in our first model, with the exception that at each reproduction event there is some probability, *q*, that instead of giving rise to another *G* cell, a different cell type, denoted *S*, is produced instead. *S* cells also consume resources, but unlike *G* cells, *S* cells cannot replicate or be dispersed. The production of *S* cells is thus costly: they deplete resources and reduce the number of cells available for dispersal. Full mathematical details of the model are given in Methods and the Supplementary Information file. The phenotype of *G* cells is quantified by their growth rate, *β*, and the probability of production of *S* cells, *q*. All other parameters are fixed.

In this formulation, *S* cells are a rough approximation for soma. Like soma, *S* cells are an evolutionary dead end. In this switch to considering *S* cells as proxy for soma, it follows that *G* cells approximate germ cells: like germ cells, these dispersing cells found the next collective generation.

To connect with our previous results, simulations of the model were first performed with dispersal depending solely on the number of *G* cells within the patch at the time of dispersal, thus patches that optimise the number of *G* cells maximise the number of descendent patches. As to be expected given the cost of maintaining *S* cells, in repeated simulations of the model in which mutation affects both growth rate and the probability of production of *S* cells, the rate of *S* cell production under both slow and fast dispersal regimes declines to zero (Figure 6A-C). The equilibrium fitness of both cells and patches tend to the same values as in the previous model.

**Figure 6.**
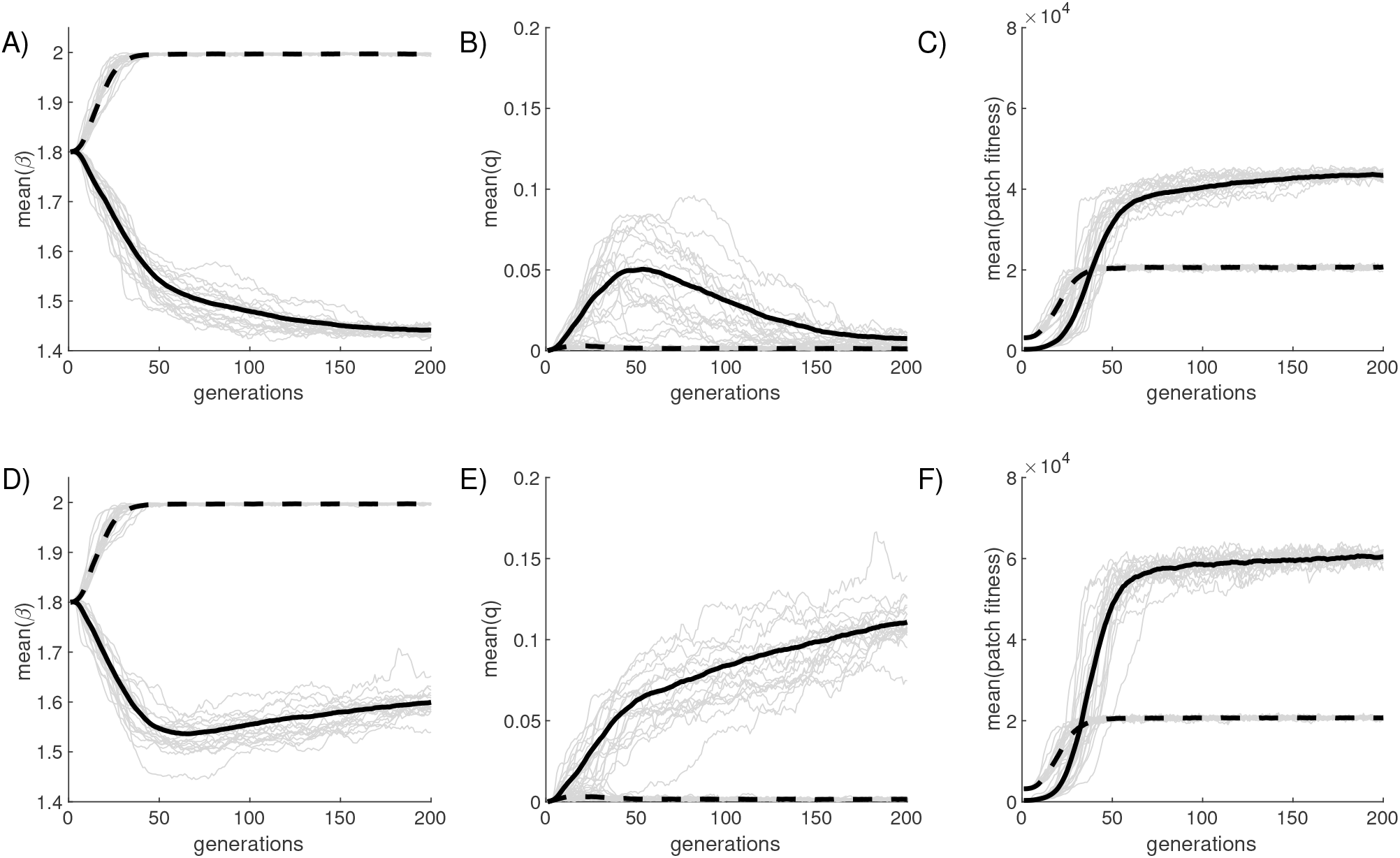
Simulations of the two-type model. A, B and C show the simulations the the model where the probability of dispersal only depends on the number of G cells in the patch at T. A shows the mean growth rate (averaged over generations), B is the mean probability of a reproduction event creating a B cell, and C shows the mean patch fitness. Each grey line is a single stochastic realisation and the solid and dashed lined are averages over 100 independent realisations for slow (T=30) and fast dispersal (T=10) respectively. Panels D, E, and F, show the similar simulations, but where the probability of dispersal from a patch is a function of both the number of G and S cells (*ρ* = 0.02). In all scenarios, the maximum cell growth rate is limited to 2. Figure S8 shows the dynamics run over 2000 generations to confirm that equilibrium is reached for the case where dispersal is slow and both cell types contribute towards dispersal.

To determine whether the ecological scaffold established by patchily distributed resources and dispersal between patches establishes conditions favouring evolutionary emergence of a division of labour, the model was re-run, but with *S* cells now endowed with ability to aid dispersal of *G* types. Mathematically this was achieved by defining the probability that a cell within a patch is chosen for dispersal be a function of both the number of *G* and *S* cells in the patch (see Methods and the Supplementary Information file for details). The strength of the additional contribution by the *S* cells to the probability of dispersal is controlled by the parameter *ρ* which is initially set at 0.02 per cell (see Methods).

As shown in Figures 6D-F (and especially 6E) *S* cells are favoured under the slow dispersal regime (the probability of *S* cell production rapidly evolves away from zero and plateaus at 0.15). Under this scenario the equilibrium cell growth rate is higher than when dispersal depends solely on the number of *G* cells (contrast this with the solid lines in Figures 6A and D). This is because increased production of *S* cells slows the rate of production of *G* cells allowing the population to peak at a comparatively higher growth rate. Mean patch fitness depends on the contribution that *S* cells make toward dispersal of *G* types.

The time to fully reach equilibrium for the case where *T* = 30 and where both cell types contribute towards dispersal, is much longer for this model than in the previous version. This is because the fitness landscape is flat in the region of the equilibrium point. This means that a broad combination of values of *β* and *q* give very similar patch fitnesses. This also manifests in large fluctuations in the value of *q* seen for individual realisations in Figure 6E. Simulations that run for 2,000 generations show eventual convergence (see Figures S7 and S8 in the Supplementary Information file). The strength of the dispersal assistance, *ρ*, affects the exact equilibrium, but over a certain threshold becomes relatively insensitive to the precise value (see Figure S10 in the supplementary material). This is because in the model there is an inherent tradeoff between the production of *S* and *G* cells.

Under the fast regime, *S* cells are not favoured and the growth rate of *G* simply increases to its maximum limit. However, if the maximum allowable growth rate increases beyond the limit of *β* = 2, production of *S* cells under the fast dispersal regime can be favoured. The key point is that under the slow dispersal regime, production of *S* is always favoured to some extent. When dispersal time is fast, production of *S* cells is favoured only if cell growth rate can increase to the point at which peak population size is reduced (through early and rapid depletion of resources). In real systems it is likely that cells would already be close to their maximum growth rate and thus further increases would depend on rare beneficial mutations. In contrast, decreases in growth rate are readily achieved via deleterious mutations.

## DISCUSSION

The major evolutionary transitions in individuality pose some of the most intriguing and complex problems in biology. Numerous perspectives have been offered, ranging from theoretical multi-level selection frameworks^3,45–50^, to views that give prominence to explanations for the evolution of cooperation^51–53^; from perspectives that emphasise the importance of specific mechanisms^4,54–56^ through those, like us, that emphasise the pivotal importance of the origins of group-level Darwinian properties^5,7,9,11,13,57,58^.

Encompassed within these diverse views are central concepts that can appear ambiguous. This is particularly true of scenarios in which ETIs are described in terms of “shifts in levels of selection”, or more specifically, shifts between multi-level selection frameworks MLS1 (where individual cells are Darwinian) and MLS2 (where groups are Darwinian). A thorough analysis lead Okasha^3^ to conclude the existence of a “grey area” between early and later stages of ETIs where both a MLS1 and MLS2 perspective can be taken. Closely allied is the notion of “fitness decoupling”^39^ — a sense that as selection shifts from a lower to a higher level the fitness of the higher level decouples from that of the lower — and the related idea of “de-Darwinisation” of lower level components^5^. While to the initiated all these terms convey meaning, they remain largely metaphorical and descriptive (but see Michod and Nedelcu^35^ for a model): discussion of issues surrounding ETIs needs to become more mechanistic.

Our mechanistic approach places emphasis on simplicity, causality and gives prominence to ecological factors. The ability of natural selection to act on collectives of cells depends on emergence of some manifestation of heritable variance in fitness at the collective level. In our take-nothing-for-granted approach the possibility that this arises from co-option of pre-existing cell-level traits was recognised, but put aside. While resulting in a high bar, it gives emphasis to the fact that reproduction, heredity and variation are derived traits and their existence should not be presumed^7,11,14,59^. It has also made transparent a genuine dilemma, namely, the need to explain how Darwinian properties emerge from non-Darwinian entities and thus by non-Darwinian means. If Darwinian properties do not pre-exist, or cannot arise by co-option of pre-existing lower-level traits, then their earliest manifestation necessarily lies in some exogenous factor(s). The solution we advocate involves recognising the continuity between organisms and their environments; the idea that Darwinian-like properties can be scaffolded by the environment in much the same way that reproduction in viruses is scaffolded by the host cell^5^, or that development can be scaffolded by overlap of parts between parents and offspring^60^.

The mechanistic models outlined here show that certain ecological structures can scaffold Darwinian-like properties on collectives, causing the constituent cells to experience selective conditions as if they were members of nascent multicellular organisms — even to the point where traits emerge that are defining features of multicellular life. The circumstances are minimal: nothing more than patchily distributed resources and a means of dispersal between patches. The existence of patches ensures that variation among collectives is discreet, while establishment of future recurrences of patches via single founding cells not only reinforces discreteness but is akin to reproduction. At the same time passage through a single cell bottleneck establishes high fidelity between parent and offspring patches. Together, this scenario combined with our mechanistic approach, captures in a single model, essential features of the transition from cells to multicellular life including the transition between MLS1 and MLS2 frameworks.

The second timescale is of critical importance in that it underpins a death-birth process at the level of patches^30,61,62^. Without this feature there would be no, or minimal, evolutionary impact of the second timescale on the fate of cells. Patches fail or succeed based on properties of the cells. The fact that slow growing cells are favoured when dispersal time is long is a direct consequence of the feedback between the patch-level birth-death process and the evolutionary dynamics of cells. Although within-patch selection favours fast growing cells, patches dominated by fast growing cells contain few viable cells for dispersal. Long-term success of cells thus comes from alignment of cell and patch fitness. The model thus explicates further the concept of fitness decoupling^35^. If the growth rate of cells is fast relative to the longer timescale, such that there is suboptimal patch occupancy at the time of dispersal, then patch-level selection will drive the evolution of reduced cell growth rate, leading to enhanced patch fitness.

Parallels exist in models of virulence evolution in pathogens^63–68^. These are evident in the use of mechanistic models, but also in the model structure and broader findings. The trade-off between growth rate and dispersal has been previously studied, but differently framed. For example, in models of the evolution of virulence, high external mortality of the host is known to favour the evolution of virulence^68^. In our model this is equivalent to the case where dispersal acts on a fast time-scale compared to peak population time. On the other hand, models on the trade-off between growth rate and growth yield have shown longevity of the host to be critical for the evolution of growth rate^69^, which echos our findings when the dispersal time is long. Given the similarity of our models and assumptions, it is likely that other findings from this field may be important for understanding the consequences of ecological scaffolding and the development of further models. In this direction, our model can be mapped directly to a multi-strain susceptible-infected-recovered (SIR) epidemic model^70,71^, where cell types correspond to infected individuals with different pathogen strains and susceptible types are analogous to within-patch resources. This opens the possibility for modelling and even formulation of predictions regarding viral division of labour^67^.

An at first unexpected, albeit important, subtlety surrounding the second timescale arises from its frequency of occurrence relative to the initial growth rate of cells. Beginning from a position where the growth rate of cells leads to suboptimal patch occupancy at the time of dispersal, as in Figure 3, a dispersal time that coincides with the exponential growth phase of cells drives an increase in cell growth rate (Figure 3C and 3D), while also marginally increasing patch fitness. More significant though is the fact that the fast dispersal regime is not conducive to the evolution of a reproductive division of labour. Under the fast dispersal regime growth rate of cells is the sole factor governing patch success: any reduction in total yield of *G* cells due to production of *S* cells is not offset by contributions that *S* cells make to dispersal.

For the evolutionary emergence of a reproductive division of labour the second timescale needs to occur when cells are not in exponential growth phase (Figure 3A and 3B). When evolutionary success depends on being the fastest, limited opportunities exist for exploration of phenotypic novelty^72–74^. This stems from the simple fact that manifestations of phenotypic novelty typically come at some cost to growth rate. In other words, opportunities for evolution to explore phenotypic space require the possibility that slow growing types not be excluded by selection. Selection comes to reward persistence and not simply types that grow fastest.

There are additional aspects of the longer timescale that are illuminating. It is usual when discussing the transition from cells to multicellular life to consider the cell the “lower level” and the group the “higher level”. Accordingly, the transition from cells to multicellular life is often referred to as a levels of selection problem^3,5,45,48^. The same is true for any of the major ETIs. It is clear from our patch model that the evolution of multicellular life might be better articulated as a problem to be solved by understanding conditions leading to the emergence of a second timescale over which a birth-death process operates on discretely packaged variation. This shift from levels to timescales does much to clarify the kinds of conditions necessary to effect ETIs^22,37,38^.

The occurrence of ecological conditions in nature that generate birth-death dynamics at a timescale longer than the doubling time of particles are likely rare, perhaps explaining why transitions to multicellular life are rare^13,75^. Elsewhere we have articulated spatially structured environments afforded by reeds in ponds about which surface-colonising microbial mats form, allowing cells to access oxygen that is otherwise limiting. Periodic collapse of mats marks death events that allow the possibility that new mats arise by dispersal from extant types^7,21^. Precisely this scenario underpinned design of on-going lineage selection experiments exploring the evolution of multicellular life^19,20^.

Once attention focusses on dispersal events, along with recognition that such events may effect collective reproduction, then attention turns to emergence of simple primordial lifecycles. Those involving more than a single phase, for example those involving soma- and germ-like phases, establish by virtue of the life cycle, a second timescale over which selection acts^19^. A significant aspect of such life cycles is the fact that birth-death events depend on the efficacy of developmental processes that become the focus of selection. Indeed, the evolution of lifecycles is intimately connected to the transition of multicellular life^9,13,76^. Even in our simple conceptual model, the moment that *S* cells have a selective advantage, then a rudimentary life cycle manifests, with selection able to set the developmental programme via effects on *q*, the rate at which *S* cells are produced. Arguably this marks an early step in the process of endogenisation: the process by which externally imposed Darwinian-like properties become integral features of the new entity^3,14^. Another possibility would be for *S* cells to provision new environments with resources as does the cotyledon in a plant seed, thus freeing evolving collectives from dependence on patchily distributed resources. In the *Pseudomonas* experiment, as generations of selection have passed, dependence of the evolving lineages on the scaffold has lessened. This is especially noticeable via evolution of a simple developmental programme controlling the switch between soma-like and germ-like phases^19^.

Thus far we have been silent on cooperation and the causes of cooperation that typically feature so prominently in discussions on the evolution of multicellular life^52,77^. There is no doubt that cooperation and integration are basic features of multicellular organisms, but the scaffolding perspective does not presuppose cooperation among cells as a necessary first step. Nonetheless behaviours (interactions) that might reasonably be labelled as cooperation stand to evolve given appropriate ecological scaffolds. For example, the slow growing cells favoured when dispersal time is slow (Fig 3A, 4A and 5A) could be labelled cooperating types because they show restraint in the face of plentiful resources and the fast growing mutants arising within patches might be termed selfish types, but there is no need to use such labels. Indeed, applying these labels brings focus to the individual cell, and detracts from the ecological context and importance of timescales that is critical to scaffolding, and to where causal processes lie.

One behaviour often labelled as an extreme form of cooperation — suicidal altruism — is evident in the *S* cells. Such behaviours can be seen from the perspective of single cells with the temptation to invoke inclusive fitness^78^, but to do so, would be to ignore the importance of ecology^79^ and population structure^17^. It is the meta population structure determined by ecological circumstances that ensures patches are founded by single cells and this both limits within patch conflict, while also being necessary for the emergence and maintenance of a reproductive division of labour. In our model within-patch close kinship is thus a consequence of environmental structure, with both group structure and kinship being particularly conducive to evolutionary transitions in individuality^80,81^.

In specifying our models, we have defined a minimal set of conditions that demonstrate the workings of ecological scaffolding. This has allowed elucidation of a number of general features as discussed above. The downside of a minimal model is the assumption of a strict set of ecological conditions. In particular, we have assumed a single cell bottleneck. Patches are also independent, with no interaction or migration of cells during the growth phase. This raises the question as to how these idealisations can be relaxed and how robust the model and process of ecological scaffolding.

Manipulations of our model that introduce effects akin to those arising from variation in dispersal time show that Darwinian properties imposed on collectives via an environment composed of patchily distributed resources and a means of between patch dispersal are rather robust to effects arising from asynchronous timing of dispersal. We have avoided exploring the effect of changes in other ecological factors because of the need for a more sophisticated model that will be reported elsewhere. Nonetheless, intuition — backed by findings from a new model — is that allowing patches to be founded by more than a single cell has two effects. Firstly, it leads to much stronger within-patch competition. Secondly, it reduces the relationship between patch phenotype (size at the time of dispersal) and traits of the cells that determine cell growth rate. As discussed above, the bottleneck is key for fitness decoupling as this allows slower growing cell types to spread. Under a slow dispersal regime, increases in bottleneck size undermine the effects of patch-level selection, thus slowing the evolutionary dynamic in a manner akin to reductions in the synchronicity of dispersal time.

The concept of ecological scaffolding has a number of applications and implications. In concluding we mention briefly three areas. The first concerns the emergence of the first self-replicating chemistries at the moment life emerges from non-living material. Recognition that Darwinian-like properties might emerge from the interplay between chemistry and environment opens the door to conceptual and experimental scenarios whereby chemistries that lack capacity for autonomous replication might begin to transition, through a process of templated production of bioactive compounds, toward a replicative process. Alkaline thermal vents appear to offer as much: these highly compartmentalised structures sit at the interface between hydrogen-rich fluids arising from the flow of heated water across serpentine, and acidic ocean waters with the possibility that carbon-dioxide is reduced to biologically active species^82,83^. Ensuing “growth” of products within the porous compartments sets in place the possibility of a replicative process. Incorporation of such ecological structures in future experimental designs may provide Darwinian ingredients that are typically absent from explorations of the chemical origins of life. The second area of relevance is infectious disease biology. To a pathogen, the eukaryotic host offers a discrete patch of resource. Pathogens that rely on transmission for long-term persistence experience selection over two timescales. Our model leads to the prediction that pathogens that passage through restrictive bottlenecks, such as HIV, are likely to have evolved more complex life histories than currently appreciated, involving, for example, a division of labour. This seems to be true of *Salmonella typhimurium* that switches stochastically between slow growing virulent and fast growing avirulent cell types, that invade the lumen, or colonise the gut, respectively. Cells that colonise the lumen, expressed virulence factors and trigger the inflammatory response, which benefits the faster growing avirulent cells in the gut. However, unlike cells in the gut, lumen colonising cells are killed by the intestinal innate immune defences and are thus, like soma, an evolutionary dead-end^84^. A division of labour is also hinted at in the case of HIV and other chronic RNA viruses in humans that appear to escape the deleterious effects of short-sighted within-host evolution over prolonged time intervals. Growing evidence suggests that this maybe attributable to establishment of germ-line-like lineages^85^.

The third example concerns application of ecological scaffolding for *in vitro* engineering of evolutionary transitions and particularly for top-down engineering of microbial communities in which communities eventually become a single symbiotic entity^21,77,86,87^. In the laboratory environment, and aided particularly by advances in micro / millifluidic technologies, it is a relatively trivial matter to confine populations and / or communities to thousands of discretised droplets that can then be subject to a death-birth process^21,88^. In a forthcoming paper we detail this process and its outcome for the evolution of interactions that build integrated communities.

## ACKNOWLEDGEMENTS

We are grateful to Joshua Wietz for critical review of the manuscript and valuable comment. We are similarly grateful to Silvia De Monte, Guilhem Doulcier, Médéric Diard and members of our respective teams for lively discussion. AJB acknowledges an ARC DECRA fellowship (DE160100690) and support from both the ARC Centre of Excellence for Mathematical and Statistical Frontiers (CoE ACEMS), and the Australian Government NHMRC Centre for Research Excellence in Policy Relevant Infectious diseases Simulation and Mathematical Modelling (CRE PRISM^2^). PB acknowledges a Macquarie University Research Fellowship and a Large Grant from the John Templeton Foundation (Grant ID 60811). PBR acknowledges generous financial support from MPG core funding and previously from the Marsden Fund Council from New Zealand Government funding, administered by the Royal Society of New Zealand.

## SUPPLEMENTARY MATERIAL

Supplementary Information

Supplementary Movie File 1

Supplementary Movie File 2

Supplementary Movie File 3

Supplementary Movie File 4

## METHODS

### Single phenotype model

We construct the simplest possible model to demonstrate the dynamics of two nested Darwinian populations. We assume a fixed population of patches each provisioned with a fixed amount of resource that is consumed by the cells in order to divide. The dynamics within each patch are independent with cells undergoing a birth/ death process for a fixed length of time, *T*, after which a dispersal event occurs leading to the colonisation of a new generation of patches (with replenished resources).

We first describe the basic birth / death dynamics within a patch (with no mutations). Cells have a mean life time that, without loss of generality is set to 1. We assume homogeneous mixing within the patch and hence mass-action like dynamics governing the rate of reproduction of cells, which each divide at a rate *β* multiplied by the proportion of resource in the patch. Under these dynamics, the number of cells and amount of resource within the patch follows modified Lotka–Volterra equations^89,90^, where there is no replenishment of the resource, i.e.,

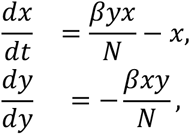

where *x*(*t*) and *y*(*t*) are the number of cells and amount of resource at time *t* respectively and *N* is the initial amount of resource in the patch. The initial conditions for these equations are *x*(0) = 1, representing the initial founding cell, and *y*(0) = *N*. The population initially grows exponentially with the resource being depleted at the same rate. At some point, the resource becomes significantly depleted leading to a reduction in growth rate. The population peaks once the rate of growth matches the rate that cells die and the population of cells declines (the trajectories shown in Figure 1 are examples of these dynamics). The total number of possible cell divisions and hence the peak population size is limited by the initial amount of resource in the patch; high growth rates lead to large populations that peak early, but which then also decrease quickly.

In the is simplest version of the model, we fix *N* = 10^6^ for all patches. To investigate how environmental predictability affects the dynamics we can introduce variability into the within-patch dynamics in two ways. We can let *N* ∼ *norm*(10^6^, *σ*_*N*_), i.e., the initial amount fo resource is sampled from a normal distribution independently for each patch in a generation. For the formulation of the model described above, where the growth rate is scaled by *N*^−1^, this amounts to assuming that the patch volume scales with the initial resources, so the concentration remains fixed. Alternatively we can introduce variability into the initial condition, *y*(0) ∼ *norm*(*N*, *σ*_B_), while keeping *N* fixed at 10^6^, which then introduces variability in the initial concentration and hence the initial growth rate.

The possibility of mutations are added to the basic model, which are assumed to only affect the growth rate of cells. For simplicity, possible growth rates are discretised (with step size *μ*) so the model tracks the populations of each type with a particular growth rate. As before, the mean life-span of all types is fixed at 1. The single cell bottleneck and limit to the amount of growth within a patch imposes a strong homogeneity on the composition of a patch. This is because mutants of the founding cell type cannot reach appreciable numbers within a single growth phase and mutants produced by mutants are correspondingly much rarer and can be ignored to a first approximation.Thus in our model we limit the number of mutant types tracked to the first two, one step higher and lower than the founders growth rate, (hence each patch has at most three strains within it). Thus if the founding cell has growth rate *β*_0_, the mutant strains will have growth rates *β*_0_ ± *μ*.

To model the dynamics of this expanded system we adopt a piecewise deterministic approach, where the times of the introduction of new types (with different growth rates) via mutations are modelled stochastically, but the growth dynamics between these times are modelled deterministically^91^. At each division event we assume a probability, *p*, of creating a child cell with a different growth rate, and hence with probability 1 − *p* a cell of the same type is produced. The times at which mutants are introduced can then be stochastically simulated as outlined in the Supplementary Material.

Between the introduction of new types, the numbers of each type already in the patch and the amount of resource evolves via a set of coupled ODEs,

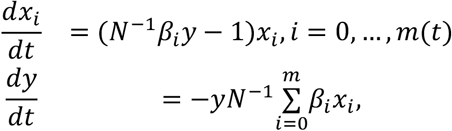

where *β*_*i*_ is the growth rate of the *i*’th cell type and *m*(*t*) is the number mutant types currently in the patch (so if *m*(*t*) = 0, only the initial colonising type is present). This is equivalent to Lotka–Volterra dynamics with competition, but where resources are not replenished. An example trajectory from this model is shown in Figure 2. The use of a piecewise deterministic model, where times of new mutants arising are stochastic but the growth dynamics are otherwise deterministic is a pragmatic comprise between computational efficiency and realism, but ignores other stochastic effects, such as the time for the cells to grow to an appreciable number before exponential growth is fully underway. Full stochastic models^31^ can account for this and display a distribution of patch sizes with a much larger variance similar to simply including variance in concentration of resource as done in this paper. Forthcoming work shows that the evolutionary dynamics are similar.

Simulation of the full model over multiple generations proceeds as follows. Each of the *M* patches are seeded with a single cell, with growth rates determined from the previous dispersal step. In the first model, all patches experience the same fixed growth time until dispersal, *T*. We can relax this by allowing growth periods for individual patches within a generation to be sampled from a normal distribution, *τ* ∼ *norm*(*T*, *σ*_*T*_), where *σ*_*T*_ is the variance and *T* is now the mean. As dispersal is taken to occur simultaneously across the patches, this variance in growth time is taken to arise in the time for the cells to be first deposited in the patches after dispersal.

For the first model, the dispersal dynamics only depend on the numbers of cells within each patch. A new generation of patches is founded by randomly selecting a patch in proportion to the number of cells within it and then randomly selecting a cell, within the chosen patch, again in proportion to its number within the patch. This procedure is equivalent to simply picking particles randomly from the whole population of patches. Hence the larger the number of a given type within the population, the more likely it is to be dispersed. This two-step procedure is simulated for a given number of generations and quantities, such as the average growth rate within a generation, can be calculated. Because the model is mechanistic, it is possible to track the genealogies of both the cells and patches, as shown in Figure S4 in the supplementary material.

### Sterile cell model

We take a similar approach to simulating this version of the model, employing a piecewise-deterministic approximation where the times of the introduction of new types due to mutations are stochastic and the birth / death dynamics are deterministic. As the *S* cells only interact with the *G* cells via competition for resources and influence dispersal when they have reached an appreciable number, the growth of the *S* types is assumed deterministic rather than stochastic. Hence between mutations, the dynamics evolve as

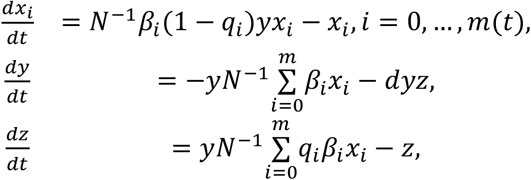

where *z*(*t*) is the number of *S* cells in the patch. The initial conditions will be ***x***(0) = (1,0,0), *y*(0) = *N*, and *z*(0) = 0. This model introduces two new parameters: *q*_*i*_, which is the per event probability of producing an *S* cell and *d*, which is the rate at which *S* cells consume the resource. The parameter *d* remains fixed, but *q*_*i*_ is subject to mutation. As the phenotype space is now two-dimensional the scheme for generating mutants is different to the first model, but the number of new mutants remains limited to the first two produced by the founding type. We assume that only *G* cells can be dispersed, hence the bottleneck enforced by the dispersal mechanism means a patch is always seeded from a single *G* type cell. More details of the simulation procedure and mutational process are given in the Supplementary material.

For dispersal we assume the system to be composed of *k* = 1, …, *K* patches. Then let 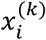 be the number of type *G*_*i*_ and *z*^(*k*)^ be the number of type *S* in patch *k*. Each patch is founded by a single *G* cell with phenotype (*β*, *q*)^(*k*)^. These cells reproduce, mutate, and create *S* cells until the dispersal time, *T*, at which point a sample is taken from the resulting populations to create the next generation. This occurs in two steps:

1. Randomly select a patch *k*, in proportion to its weight, *w*_*k*_, which is function of the proportion of its constituents, 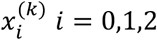 and *z*^(*k*)^.
2. From the patch selected in step 1, randomly select a G cell from the total patch population.

To assign the weigh to patches in step 1, we use the function,

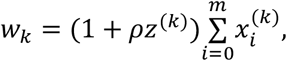

where *ρ* is the additional benefit to the dispersal process per *S* cell in the patch. This can be interpreted as the *S* cells aiding dispersal from the patch, for example by attracting the dispersal agent. If *ρ* = 0, then the dispersal process is as in the first model, i.e., the probability of choosing a patch is proportional only to the number of *G* cells, so patches with more *G* cells at the time of dispersal are more likely to be sampled from.

