## Supplementary Information for "Ecological scaffolding and the evolution of individuality: the transition from cells to multicellular life"

September 25, 2019

#### 1 Single phenotype model

##### 1.1 Piecewise deterministic approximation

Our model is a piecewise deterministic process where mutations that create new types occur as an inhomogeneous Poisson process [1] with rate

$$\lambda(t) = p\,N^{-1}y(t)\sum_{i=0}^{m}\beta_i x_i(t), \tag{1}$$

where $x_i(t)$ is the number of cell of type $i$ at time $t$, with corresponding growth rate $\beta_i$, and $y(t)$ is the amount of resource at time $t$. The sum in (1) will be dominated by the term from the initial cells that colonise the patch. Between mutations, the dynamics evolve as a set of ODEs,

$$\begin{aligned} \frac{dy}{dt} &= -yN^{-1}\sum_{i=0}^{m}\beta_i x_i \\ \frac{dx_i}{dt} &= \beta_i N^{-1} y x_i - \gamma x_i \quad i = 0,\dots,m(t). \end{aligned} \tag{2}$$

Note that we have neglected a term of $1-p$ in the Eqs (2) as this is close to 1. As each patch is colonised by a single cell, the initial conditions are $m=0$, $x_0(0)=1$ and $y(0)=N=10^6$. When a new mutant type is created, the state of the process (and hence the number of of ODEs to be solved) is updated as,

$$\begin{aligned} m &\leftarrow m+1, \\ y(t) &\leftarrow y(t)-1, \\ x_m(t) &= 1. \end{aligned} \tag{3}$$

This model can be derived in a number of different ways from an fully stochastic model and different assumptions result in slightly different formulations of the above equations. Here we have chosen the simplest, so we do not include the effect of later mutations between already existing types in the ODEs (2). We also limit the the number of mutant types to 2 created by the founding type (with growth rates $\beta_0 \pm \mu$) and ignore back mutations, which will be limited anyway by the small numbers of mutants produced overall due to the bottleneck and patch ecology. The probability of a faster or slower growing mutant being produced first is set at 1/2 for simplicity.

To solve for the dynamics of a patch we need to determine the mutation times, i.e., we need to sample from the inhomogeneous Poisson process with rate given by Eq. (1). There are a number of possible ways of doing this, but we use a cumulative distribution function (CDF) inversion method because, as shown below, this allows the use of basic ODE solvers for the whole process. The expectation of the Poisson random variable with rate (1) is

$$E[N_t] = \Lambda(t) = \int_0^t \lambda(s)ds.$$

It can be shown that the inter event time, $X_j$, conditioned on the first $j$ times $t_1, \ldots, t_j$ has CDF [7, 9],

$$F_{t_i}(x) = 1 - \exp(-\Lambda(t_i + x) + \Lambda(t_i)). \tag{4}$$

To use a CDF-inverse method to generate the times from (4) we then need to solve the equation [3]

$$\int_t^{t+x} \lambda(y)dy = -\ln(u)$$

where $u \sim \mathrm{U}(0, 1)$ is a uniform random variable, $t$ is the current time and $x$ the time until the next event. Using equations (1) and (2) we see that

$$\lambda(t) = -p\frac{dy}{dt}. \tag{5}$$

hence

$$-Np\int_t^{t+x} \frac{dy}{dt}dy = -p(y(t+x) - y(t)) = \ln(u). \tag{6}$$

Hence Equation (5) can be solved simultaneously with the equations (2). The procedure is to first generate $u$, then starting with the initial condition we integrate the combined set of equations (2) and (5) until equation (6) is true, or the dispersal time, $\tau$, is reached. This is easily accomplished using an event function passed to the ODE solver. In this paper we use MATLAB's built in function `ODE45`. Once the time has been determined the state is updated according to Eq. (3).

#### 1.2 Dynamics

Figure S1 shows four realisations of the within patch model with different founding cell growth rates. The final panel shows the trajectory resulting from the optimal growth rate for the dispersal time of $T = 30$, but note that the peak does not coincide with $t = 30$, but occurs slightly before. This figure also illustrates that slower growth rates lead to smaller *peak* populations in an absolute sense.

Another way to interpret the evolutionary dynamics of this model is by thinking in terms of a fitness landscape. This is possible because the accumulation of mutants in the patch is small, and so the number of cells at the time of dispersal is strongly determined by the growth rate of the founding cell. A fitness landscape is derived by plotting the patch size (number of $G$ cells at the time of dispersal) as a function of the growth rate for different (fixed) dispersal times with $p = 0$, i.e., no mutation. Figure S2 shows this for $T = 10, 15$ and 30. As patch fitness is proportional to patch size, larger patches at the time of dispersal will be more likely to be selected. For a given $T$ the system will evolve up the fitness curve; for example if $\beta = 1.8$ initially and $T = 30$ (blue curve), then this predicts that the average $\beta$ in the system will decrease, as this leads to larger patches. Conversely, if $T = 10$ then the growth rate increases. The dashed line shows the imposed maximum growth rate, and hence if $T = 10$, this is where evolution will stop. If the maximum growth rate was unbounded then the growth rate would continue to rise, reaching an equilibrium around 2.5. By choosing a slightly slower dispersal time, $T = 15$, the equilibrium growth rate is reached at $\beta < 2$.

This fitness landscape view also emphasizes that whether we observe fitness decoupling or not depends on the dispersal time scale *relative* to the initial cell growth rate. It also shows that shorter times allow for fitter patches in an absolute sense as larger cell growth rates lead to bigger overall sizes. This is simply due to the particular form of the birth / death dynamics assumed in the model. Figure S3 shows a genealogical representation of the evolutionary dynamics, discussed more in the main text.

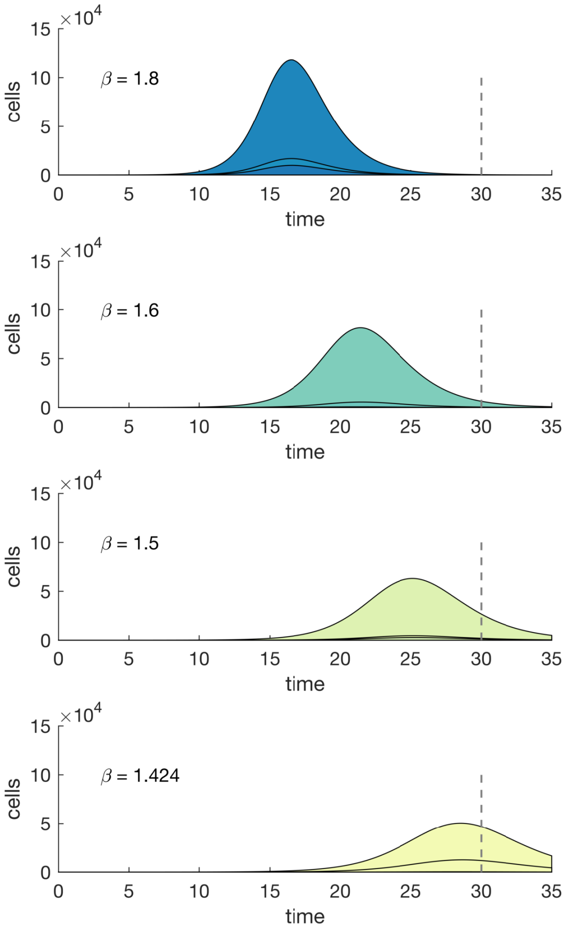


Figure S1: Realisations of within-patch dynamics with mutations for different initial cell growth rates. $\beta = 1.42$ is the optimal growth rate for $T = 30$. The curves are colour filled according to their growth rates as in the main text (darker blues represent faster rates and lighter greens slower rates). Other parameters: $N = 10^6$, $p = 10^{-2}$ and $\mu = 0.02$.

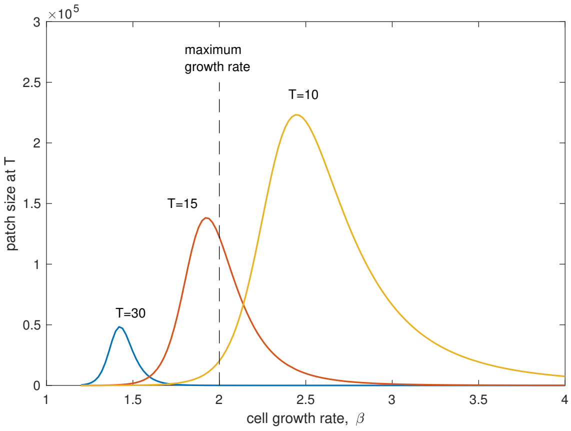


Figure S2: Fitness landscape for the single type model with no mutations. For the indicated dispersal times ($T = 10$, yellow; $T = 15$, red; and $T = 30$, blue) the patch size (population at the dispersal time) is plotted against the initial cell growth rate. The maximum possible growth rate that is enforced in our models is shown by the dashed lined.

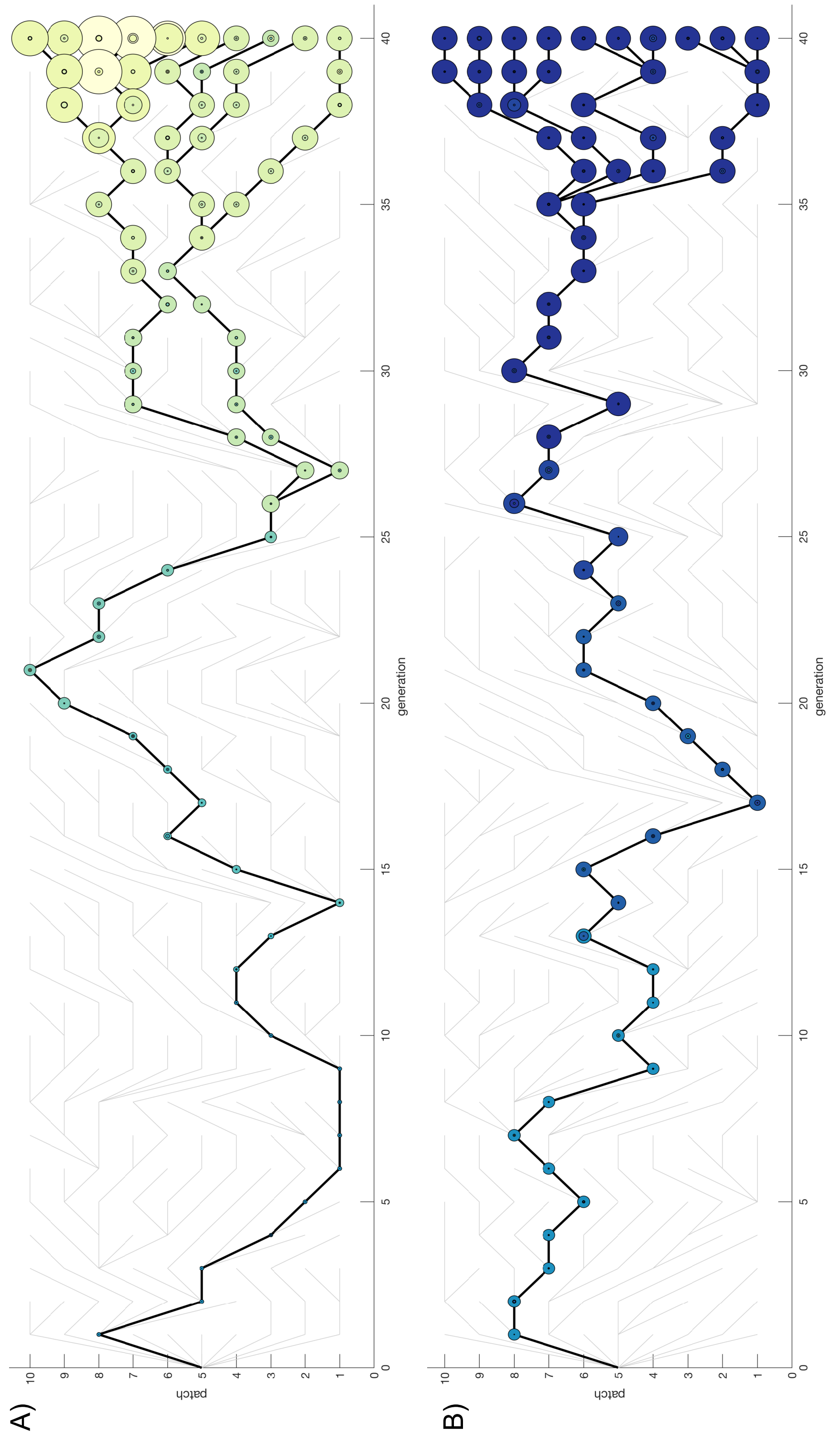


Figure S3: Genealogy of patches under slow (A) and fast (B) dispersal regimes. The simulations to produce these have only 10 patches and modified mutational parameters compared with those in Figure 2 of the main text. This is to allow a clearer visualisation of the process, which otherwise requires many more generations to see change. Video versions of these are also included in the supplementary material. As in figures 4 and 5 of the main text, the cell numbers in each patch are proportional to the area of the circles and the growth rates are indicated by the colours, as shown by the colour bar in figure 4 of the main text. The mutational parameters are larger for these simulations ($\mu = 0.05$, $p = 0.05$) so evolution occurs on a quicker timescale as compared with the results shown in Figure 3 of the main text.

### 2 Additional asynchronous dispersal figures

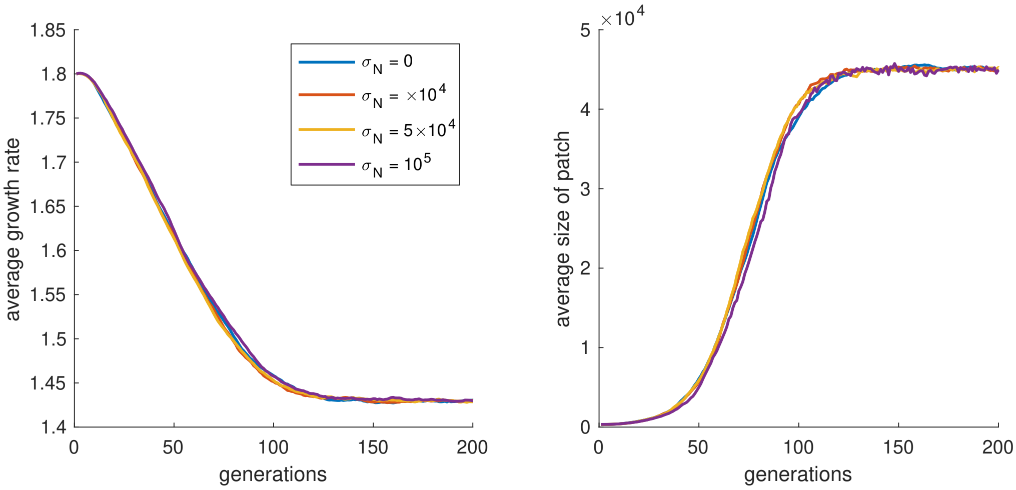


Figure S4: Evolution of the average cell growth rate and patch size with increasing variance in the initial resource within a patch. Each line is calculated from the average of 50 independent simulations. Variance is introduced by sampling $N \sim \mathrm{norm}(10^6, \sigma_N)$ for each patch in each generation. The initial condition is set as $y(0) = N$, so in effect this means the *concentration* of resource within the patch remains fixed. The variability in the resource then leads to variability in the sizes of the patches at the time of dispersal, but not in their time to peak. Hence the evolutionary dynamics do not change.

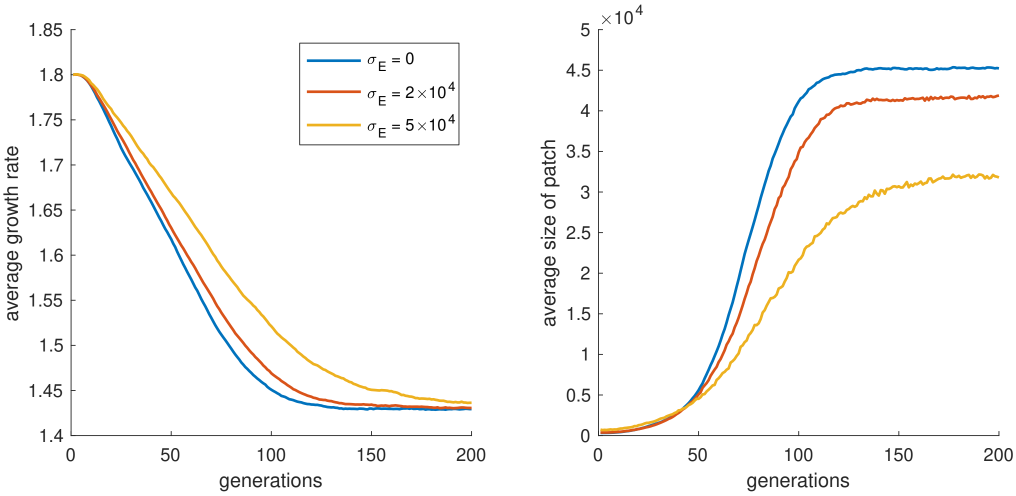


Figure S5: Evolution of the average cell growth rate and patch size with increasing variance in the initial concentration of resource within a patch. Each line is calculated from the average of 50 independent simulations. Variance is introduced by sampling $y(0) \sim \mathrm{norm}(N, \sigma_E)$, and keeping $N = 10^6$ fixed. In this version of the dynamics, the variability in the concentration leads to variability in the time for the population to peak within a patch. Thus the results are qualitatively similar to those described in the main text, where increasing variance in the initial seeding time leads to a slowing of the evolutionary dynamics and a decrease in the equilibrium patch size.

### 3 Model with sterile types

In this version of the model, mutations to create new types occur as an inhomogeneous Poisson process with rate

$$\lambda(t) = pN^{-1}y(t)\sum_{i=0}^{m}(1-q_i)\beta_i x_i(t). \tag{7}$$

As before the sum will be dominated by the $i = 0$ term of the founding cell type. Between mutations, the dynamics evolve as,

$$\begin{aligned}
\frac{dx_i}{dt} &= \beta_i N^{-1}(1-q_i)yx_i - x_i, \quad i = 0, \ldots, m(t), \\
\frac{dy}{dt} &= -yN^{-1}\sum_{i=0}^{m}\beta_i x_i - dyz, \\
\frac{dz}{dt} &= yN^{-1}\sum_{i=0}^{m} q_i\beta_i x_i - z.
\end{aligned} \tag{8}$$

As $p \ll 1$, we have dropped the $1 - p$ term that should appear in the second equation.

To facilitate solving for the mutation times, we add another equations for the time derivative of $\lambda(t)$ in terms of the other state variables,

$$\frac{d\lambda}{dt} = py\sum_{i=0}^{m}\beta_i(q_i - 1)x_i = \sum_{i=0}^{m}\left(\frac{dx_i}{dt} + x_i\right). \tag{9}$$

The mutation times are then generated in the same way as described in Section 1.1. As the phenotype space is two-dimensional, the mutation process is implemented differently to the first model. When new mutant types are created, the mutant phenotype, $(\beta_j, q_j)$, is related to the original, $(\beta_i, q_i)$, by

$$\begin{aligned}
\beta_j &= \beta_i + 2\beta_r(u_1 - 0.5), \\
q_j &= q_i + 2q_r(u_2 - 0.5),
\end{aligned} \tag{10}$$

where $u_1, u_2 \sim U(0,1)$ and $\beta_r$ and $q_r$ specify the magnitude of mutation. Throughout we set $\beta_r = 0.05$ and $q_r = 0.02$. When calculating the mutant type we also enforce that $0 \leq \beta_j < 2$ and $0 \leq q_j < 1$, where the upper limit on $\beta$ is assumed due to physical constraints on the growth process. If a mutant is created with a phenotype outside these bounds then the new parameters are truncated to fall within the bounds above.

In the dispersal phase, patches are now sampled in proportion to their weight, which is given by

$$w = (1 + \rho z)\sum_{i=0}^{m} x_i. \tag{11}$$

#### 3.1 Dynamics

To understand the effect of the production of $S$ cells on the overall dynamics, trajectories are shown in Figure S6 without mutation ($p = 0$) for two different values of $q$. We see that $q > 0$ leads to the production of $S$ cells, but that the the number of $G$ cells peaks at a later time and at a lower overall number. This is because production of $S$ cells decreases the effective growth rate of $G$ cells as well as consuming the resource used for growth (without reproducing themselves) so there is less for $G$ cells to use for reproduction. If dispersal time is long compared with peak time, increasing $q$ causes an increase in patch fitness by both increasing the number of $G$ and $S$ at the time of dispersal.

As with the previous model, because the number of mutant $G$ cells in a patch is kept small by the bottleneck, we can adopt a fitness landscape approach for understanding the evolutionary

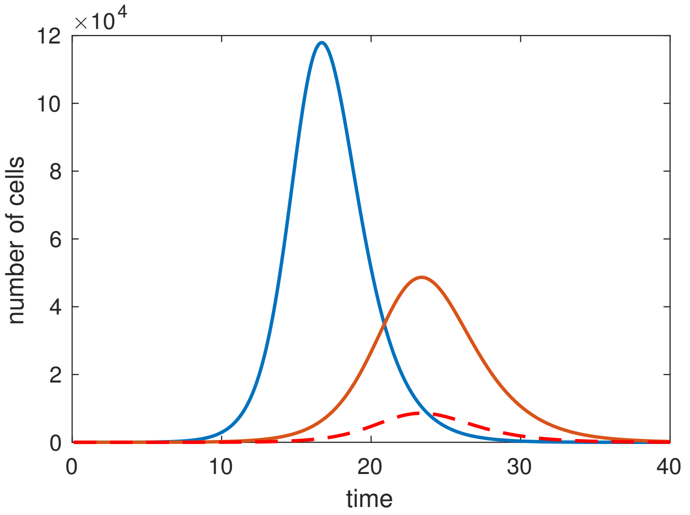


Figure S6: Trajectories from the model with sterile cells with $p = 0$. The blue line is the number of $G$ cells with $\beta = 1.8$ and $q = 0$, hence no S cells are produced. The red lines show the number of $G$ cells (solid) and S cells (dashed) for parameters $\beta = 1.8$ and $q = 0.15$. Other parameters: $N = 10^6$, $d = 2$ and $\gamma = 1$.

dynamics. This is analogous to that shown in Figure S2, but now two-dimensional. Figure S7A and B show contour plots for the number of $S$ and $G$ in a patch at the time of dispersal (assumed slow, $T = 30$) as a function of the initial growth rate, $\beta$, and $q$. Figure S7C also shows the patch fitness calculated from the weight in Eq. (11). This shows that, as expected, the number of $G$ cells is maximised by a growth rate of 1.45 and $q = 0$. Thus if patches are selected based only on the number of $G$ cells, then this is the evolutionary outcome as seen in the simulations presented in the main text. In contrast to the behaviour shown in (A), the number of $S$ cells is always maximised by increasing both $q$ and $\beta$. If the patch fitness is a function of these two numbers then this is maximised at intermediate values (indicated by $*$). Also note that the landscape is very flat in the direction of $q$ around the maximum, which is why fluctuations about the steady state seen in the simulations are large and the time to reach equilibrium is much longer for his model than the simpler version (see Figure S8).

Figure S9 shows the same quantities as Figure S7, but for fast dispersal ($T = 10$). The behaviour of the number of $G$ and $S$ cells follows a similar pattern to before, but at larger values of $\beta$. Figure S9C shows that if we impose a maximum growth rate on cells then patch fitness is maximised by maximising $\beta$ and minimising $q$ ($*$), but if no such maximum is imposed then a non-zero $q$ and larger $\beta$ maximises patch fitness ($*$). This is analogous to the behaviour shown in Figure S2, but in a two-dimensional setting. Figure S10 shows the position of the equilibrium as a function of $\rho$, and shows that over a certain value, the position is mainly determined by the inherent trade-off in the production of $G$ and $S$ cells rather than the absolute value of $\rho$.

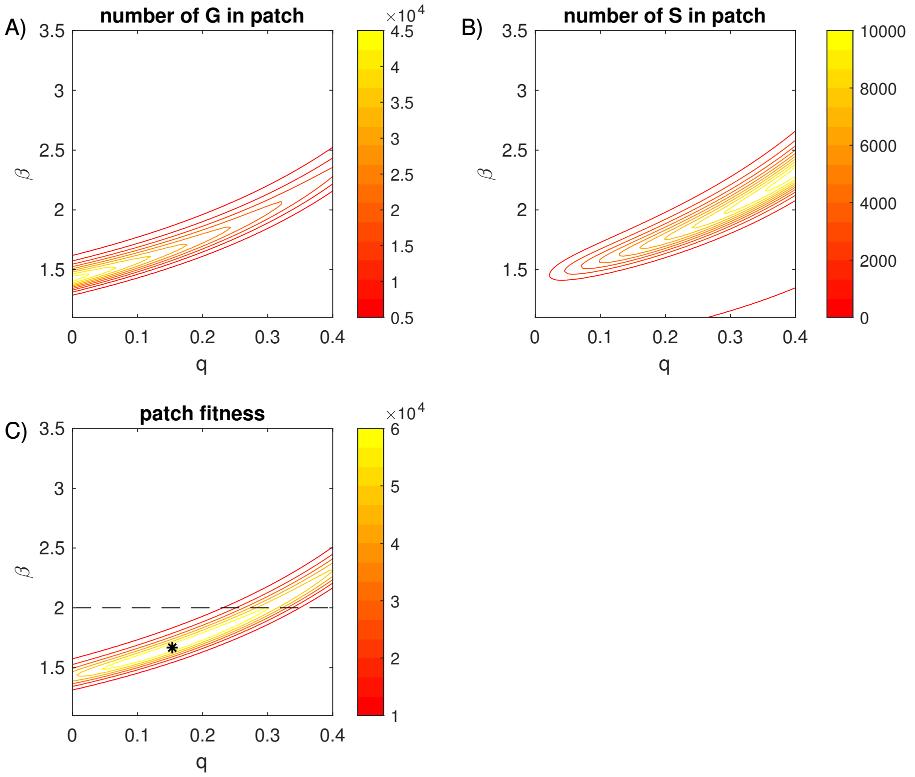


Figure S7: Fitness landscapes assuming slow dispersal ($T = 30$). (A) shows the number of $G$ cells in a patch at the time of dispersal as a function of the initial growth rate, $\beta$, and $q$. (B) shows how the number of $S$ cells at the time of dispersal changes as a function of $\beta$ and $q$. (C) shows the patch fitness calculated from Eq. (11). For slow dispersal we see this is maximised at $q = 0.15$, $\beta = 1.66$ ($*$). Other parameters: $N = 10^6$, $d = 2$ and $\gamma = 1$.

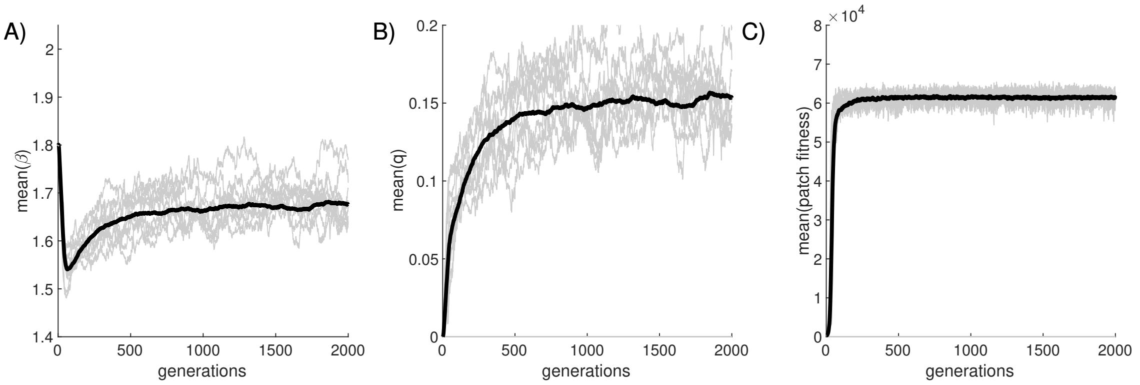


Figure S8: Simulations of the model with sterile types over 2000 generations show convergence to equilibrium as indicated in Figure S7. The comparative slowness of this convergence, as well as the large fluctuations in the mean value of $q$ for single realisations, can be attributed to the flatness of the fitness landscape about the equilibrium as seen in Figure S7C.

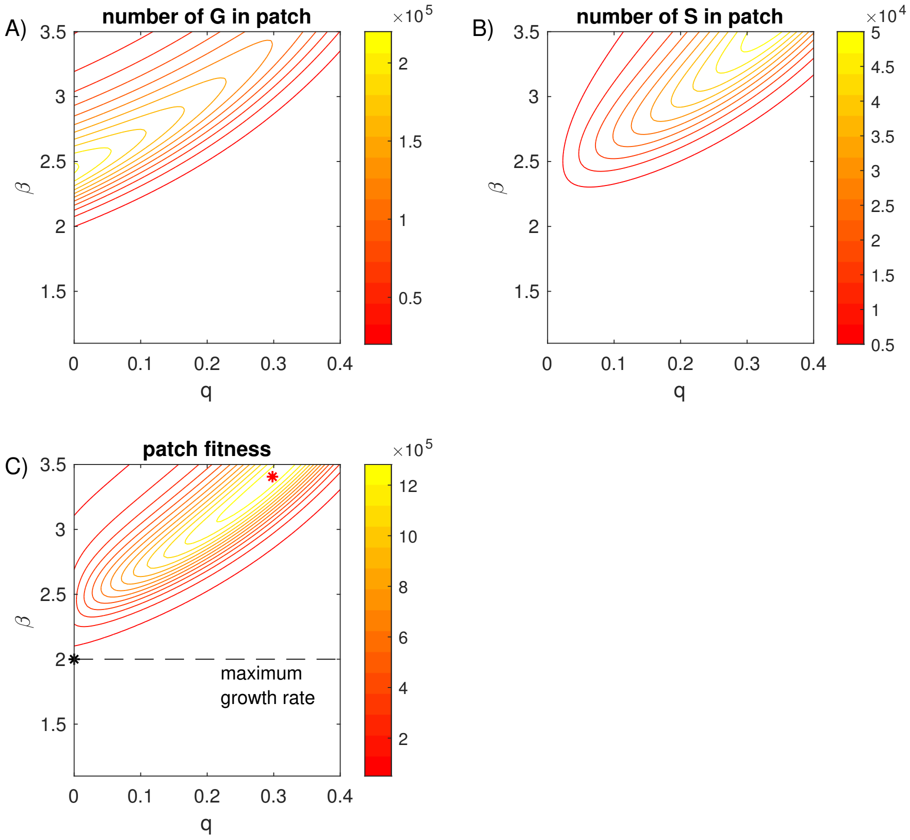


Figure S9: Fitness landscapes assuming fast dispersal ($T = 10$). This shows the same quantities as the Figure S7. If $\beta$ is limited to a maximum of 2 (shown by the dashed line) then patch fitness is maximised at $\beta = 2$ and $q = 0$ ($*$) as seen in the simulations of the evolutionary model shown in the main text. If there was no limit to the growth rate then we would see evolution to a state with non-zero $q = 0.3$ and $\beta = 3.4$ ($*$).

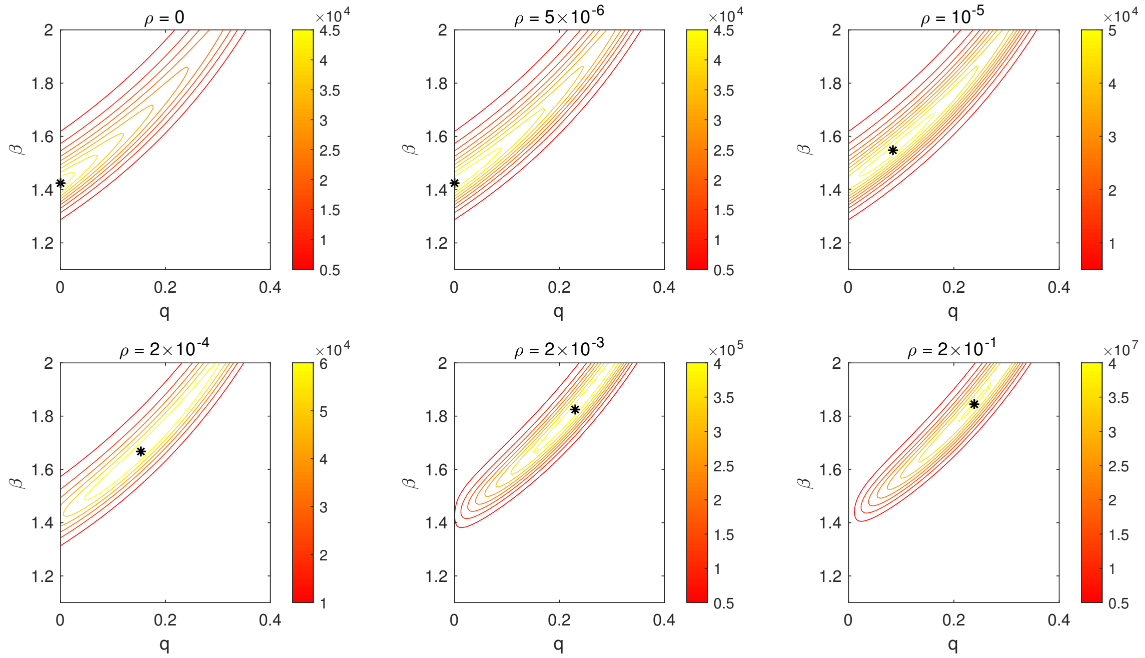


Figure S10: Change in the position of the equilibrium, indicated by '$*$' as a function of $\rho$ for slow dispersal ($T = 30$).
